# Global Landscape of Human Kinase Motifs in Viral Proteomes

**DOI:** 10.1101/2025.06.02.657064

**Authors:** Kareem Alba, Declan M. Winters, Sara K. Makanani, Prashant Kaushal, Yennifer Delgado, Immy A. Ashley, Shipra Sharma, Sophie F. Blanc, Erin Kim, Tomer M. Yaron-Barir, Jared L. Johnson, Justin Selby, Min Hur, Jocelyn Ni, Jasmine Nguyen, Matthew Lawson, Evan T. Wynn, Sophie Lin, Marc H. Brent, Elaine Yip, Ahmad Kassem, Charles O. Olwal, James Wohlschlegel, Oliver I. Fregoso, Ting-Ting Wu, Melody M.H. Li, Mehdi Bouhaddou

**Author notes:** These authors contributed equally to the work (Kareem Alba, Declan M. Winters).

## Abstract

Viruses are classically viewed as targets of host sensing, yet whether they also sense and respond to host cues remains largely unexplored. We propose that host-driven post-translational modification of viral proteins allows viruses to dynamically sense host cellular states. We annotated human kinase motifs in 1,505 viral proteomes and discovered an enrichment for stress, inflammation, and cell-cycle kinases. Mapping kinase motifs onto 21,606 viral protein structures and integrating with phosphoproteomics of infected cells revealed surface-accessible residues were preferentially phosphorylated, showed greater kinase specificity, and were under positive selection for stress and immune kinase motifs. Temporal phosphoproteomics of alphavirus-infected cells confirmed stress kinase activation and viral protein phosphorylation, and MAP kinase inhibition reduced alphavirus replication and phosphorylation of ERK and JNK motifs on viral proteins. Our findings suggest that viruses evolved as biosensors of the host signaling state, unveiling new antiviral opportunities aimed at disrupting virus decision-making.

## INTRODUCTION

Viruses must navigate a dynamic intracellular environment to successfully establish infection, replicate, and transmit among their hosts. It is well established that viruses evolved offensive strategies, such as innate immune antagonists, and hosts developed countermeasures, such as the inflammatory response, to detect and neutralize viruses.^1^ However, the extent to which viruses sense and respond to host cues in real-time, and how sensing is biochemically orchestrated, remains unclear.^2^ The capacity for rapid and context-dependent adaptation resembles a form of molecular "decision-making," enabling the tuning of viral life cycle transitions—such as replication, assembly, or exit—and maximizing viral fitness and the ability to spread amongst a host population.^3^ Furthermore, uncovering how viruses sense and respond to host environments may expose critical life cycle bottlenecks and reveal novel targets for a new class of antivirals aimed at disrupting host cues.

Viral infection is known to provoke dramatic remodeling of host signaling via post-translational modifications (PTMs).^4^ One PTM, phosphorylation, is regulated by protein kinases and phosphatases that add and remove phosphate groups at phosphoacceptor (i.e. serine, threonine, or tyrosines) residues. Phosphorylation can act as a “switch” by generating functionally distinct proteoforms in response to cellular cues.^5^ Viral proteins are no exception; phosphorylation by host kinases can expand the functional repertoire of viral proteins by modulating subcellular localization, protein-protein interactions, or enzymatic activity.^6^

Previous work revealed phosphorylation of viral proteins by host kinases plays a key role in regulating viral life cycles.^7^ During HIV-1 infection, host CDK2 phosphorylates the viral protein Tat at serine 16, enhancing transcription during S-phase and linking replication to host DNA synthesis.^8^ In severe acute respiratory syndrome coronavirus 2 (SARS-CoV-2), the nucleocapsid (N) protein exists in two functionally distinct forms, one that is phosphorylated and promotes translation, and one that is unphosphorylated and drives viral assembly.^9^ In influenza virus, phosphorylation of the nonstructural protein 1 (NS1) by casein kinase 2 (CK2) enhances its binding to double-stranded RNA, allowing it to sequester RNA and block activation of host antiviral responses.^10^ Together, these examples highlight how viral protein phosphorylation enables viruses to coordinate their life cycles with host signals.

Technological advances in mass spectrometry (MS), artificial intelligence, and genetic manipulation enable a deeper dive into how phosphorylation influences protein function and cellular phenotypes. For example, data-independent acquisition mass spectrometry (DIA-MS) has enabled unprecedented breadth and depth of exploration into the phosphoproteome.^11^ In parallel, improvements in artificial intelligence–based structural modeling using AlphaFold^12^ have made it possible to accurately predict and interrogate viral protein structures.^13^ Furthermore, recent advances in kinase motif prediction, such as the Kinase Library^14^, which integrates positional scanning peptide array data with computational prediction tools, allow researchers to assess kinase specificity for any given protein sequence. The integration of these technologies support the investigation of host kinase-mediated phosphorylation of viral proteins by enabling both large-scale detection by MS, structural interrogation by AlphaFold, and sequence-based kinase motif prediction by the Kinase Library.

In this study, we harnessed these tools to perform a global analysis of human kinase motifs across thousands of viral proteomes. By integrating kinase motif predictions for viral proteins, phosphoproteomics of infected cells, and viral protein structural modeling, we identified conserved signaling pathways that viruses can sense through exposed surface residues in their proteins. These phosphorylation-sensitive protein surfaces were conserved across diverse viral families. Additionally, we incorporated positive selection analysis to examine how kinase motifs are shaped by evolutionary pressures across viral proteomes. We further validated the functional significance of this sensing strategy by perturbing MAPK signaling during Sindbis virus (SINV) infection, revealing defects in both viral protein phosphorylation and replication. Together, our findings illuminate a previously underappreciated layer of virus-host interaction in which human kinase activity is co-opted, not just defensively by the host, but offensively by viruses to interpret and manipulate their environment.

## RESULTS

### A diverse array of viruses encode high confidence human kinase motifs

To investigate the prevalence of human kinase motifs in viral proteins, we assembled 125 proteomes of human-infecting viruses and subjected each phosphorylation acceptor residue (i.e., serine, threonine, or tyrosine), and the surrounding seven upstream and downstream amino acids, referred to as a “peptide”, to computational kinase motif analysis (Table S1). Specifically, we employed The Kinase Library^14,15^ algorithm, a recently developed algorithm informed by positional scanning peptide array (PSPA) analysis (Fig. 1A). The Kinase Library provides a score for every peptide-kinase pair; a score of 0 indicates a random association, a positive score indicates a favorable interaction, and a negative score indicates an unfavorable interaction (i.e., negative selection). The 125 viruses spanned 26 viral families (Fig. 1B), all seven Baltimore classifications (Fig. 1C), with a slight bias for RNA versus DNA-based genomes (Fig. 1D). Viral and human proteins revealed equivalent median kinase scores, which trended negative, thought to indicate a trend towards negative selection away from phosphorylation for both organisms (Fig. 1E), consistent with prior findings.^14^ A percentile score was used to compare kinase favorability for each peptide relative to >80,000 known human phosphorylation sites.^14^ We found the positive kinase score distributions to be similar in terms of their percentile ranking across viral and human proteins and used this to define a “high-confidence kinase motif” as possessing a score greater than 2, which we found exceeds the 95% percentile (Fig. 1F).

**Figure 1.**
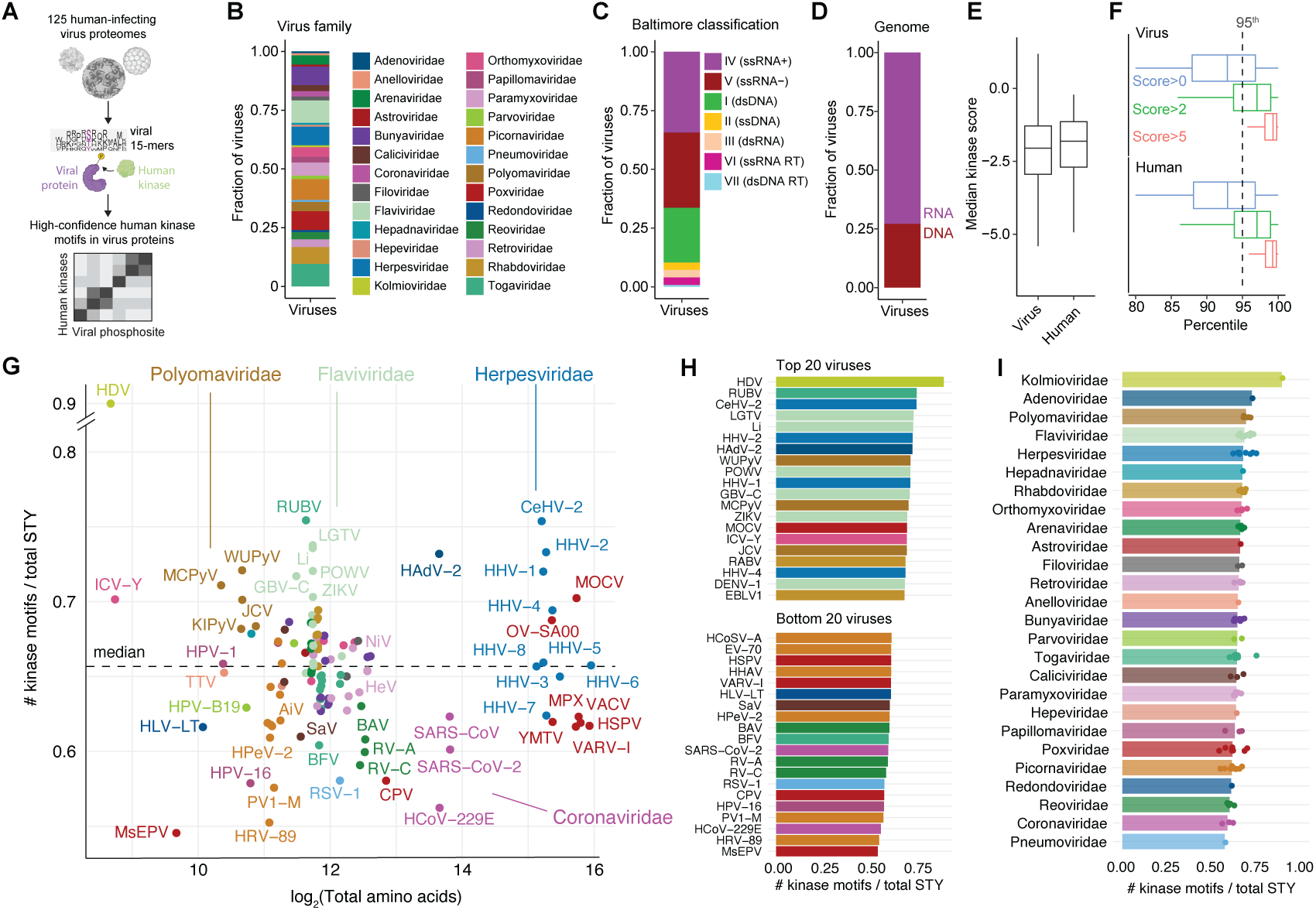
Viral proteins encode human kinase motifs. (A) A total of 125 viral proteomes were collected, and kinase specificity for various motifs was predicted using the Kinase Library algorithm. (B) The distribution of viral families represented in the dataset. (C) The distribution of Baltimore classifications among the viral proteomes. (D) The distribution of genomic material types present in the dataset. (E) A comparison of the median kinase motif scores between viral and human proteomes. (F) The distribution of kinase motif scores across viral and human proteomes. (G) The relationship between the log₂-transformed proteome size and the number of high-confidence motifs, normalized to the number of phosphoacceptor sites. (H) The top 20 and bottom 20 viruses ranked by the number of kinase motifs normalized to phosphoacceptor count. (I) The number of kinase motifs per phosphoacceptor site grouped by viral family.

We then ranked viruses by the number of high confidence kinase motifs (i.e., score > 2) divided by the number of phosphoacceptor residues (i.e., S,T, or Y) in their proteomes. We found most viruses varied in their proportion of high-confidence motifs between approximately 0.55 to 0.75, with a median of 0.66, which remained consistent for a variety of proteome sizes (Fig. 1G). Strikingly, we found Hepatitis D virus (i.e., delta hepatitis) to have the highest proportion (0.89) of high-confidence motifs for GSK3A, PIM1, and HIPK4 kinases, suggesting a role for these kinases in regulating its life cycle. In general, viruses in the *Herpesviridae*, *Flaviviridae*, and *Polyomaviridae* families had the highest proportion of high-confidence motifs, suggesting these viruses may have evolved to sense and respond to phosphorylation signaling cascades on many of their proteins (Fig. 1G-I). Conversely, some viruses possessed a low proportion of high-confidence motifs, such as the *Coronaviridae*, *Reoviridae*, and *Picornaviridae* families (Fig. 1G-I). Importantly, a virus with few high-confidence motifs may still possess functionally important phosphorylation events for key regulatory roles in its replicative cycle.

### Viral proteins encode human kinase motifs for stress, inflammation, and cell cycle pathways

To identify the kinase signaling pathways implicated in viral protein phosphorylation, we developed a statistical pipeline to compare the prevalence of kinase motif scores (KMS) between viral and human proteins, designed to identify kinases with an enriched preference for viral relative to human substrates (Fig. 2A). We reasoned that this approach would allow quantitative assessment of viral enrichment relative to the expected distribution across the human proteome. Specifically, for each kinase, we performed a Z-test comparing the KMS distribution of viral substrates (per virus) to that of all human substrates, yielding a z-score and p-value for each virus-kinase pair. A positive virus-kinase z-score indicates that a kinase preferentially targets a given viral proteome, whereas a negative z-score suggests reduced targeting, potentially reflecting selection against phosphorylation. Thus, this analysis identifies kinases that exhibit a preference (or aversion) for phosphorylating proteins within a specific viral proteome beyond the level observed in the human proteome. Using these z-scores and p-values, we constructed a high-confidence virus-kinase interaction network, representing an undirected network linking each virus to associated human kinases. For each virus-kinase pair, we applied three criteria: (i) a z-score greater than zero (z > 0; i.e., virus > human), (ii) a p-value less than 0.05 (p < 0.05), (iii) and a minimum of seven high-confidence kinase motifs (high-confidence defined as KMS > 2; Fig. S1A) detected within each viral proteome (Fig. 2A; Table S2). A threshold of seven high-confidence motifs was chosen because it yielded the highest number of enriched Gene Ontology Biological Process terms across all viruses, suggesting this threshold optimally balances the sensitivity for statistical enrichment (Fig. S1A).

**Figure 2.**
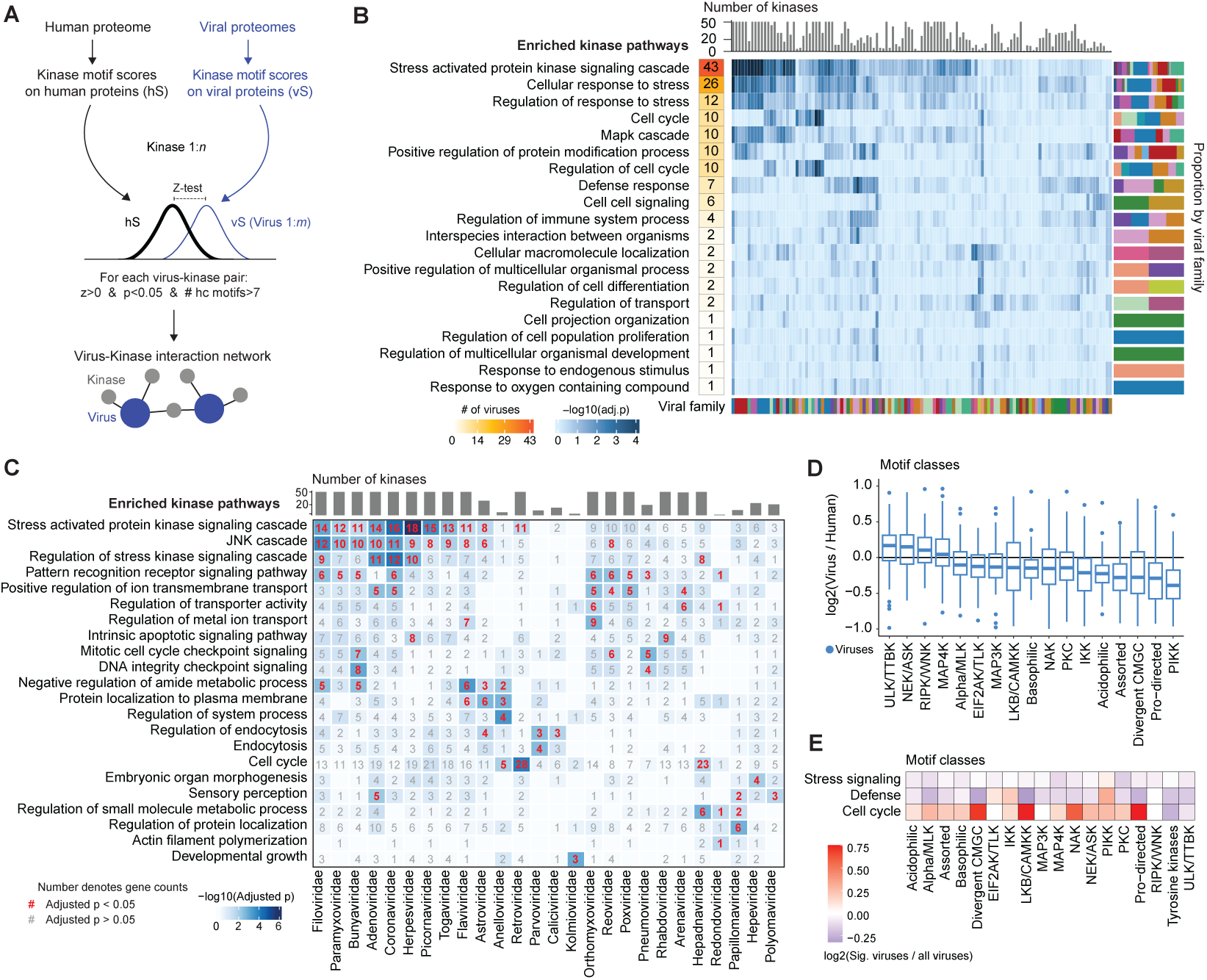
Viral kinase motifs are largely responsive to stress and inflammation kinase pathways. (A) Viral kinase motif scores were compared per virus to the global human kinase motif score distribution for each kinase, generating a z-score and associated p-value. The resulting network was thresholded based on the following criteria: (1) z > 0, (2) p < 0.05, and (3) more than seven high-confidence kinase motif scores (KMS;score > 2) per virus– kinase interaction. (B) GSOA results for top 50 high-confidence kinase-virus interactions for each virus (or all available if less than 50), sorted by the number of significant viruses per term. A list of all human kinases were used as the universe. (C) GSOA results performed per viral family. (D) For each virus, the proportion of high-confidence motifs for each kinase family was divided by the corresponding proportion in humans. (E) For viruses enriched in GSOA terms related to stress signaling, defense, and cell cycle, the proportion of high-confidence motifs was calculated, and this proportion was normalized to the baseline ratio observed across all viruses.

After identifying high-confidence virus-kinase interactions, we performed gene set overrepresentation analysis (GSOA) for each virus, using either the top 50 kinases or all available kinases if fewer than 50, to uncover the biological pathways associated with the corresponding human kinases (Table S3). The top 20 most commonly enriched terms across viruses included stress signaling cascades, MAP kinase signaling, inflammatory responses, cell cycle, protein localization, cellular differentiation, and metabolism (Fig. 2B). Interestingly, across all viruses, the most enriched pathways involved stress, inflammation, and cell cycle.

We then grouped viruses by family and repeated the Z-test analysis described above, this time omitting the high-confidence kinase motif filtering step, followed by the GSOA. This analysis revealed similar pathways as the virus-specific analysis, including stress, inflammation, cell cycle, development, and protein trafficking, among other pathways (Fig. 2C). Notably, *Herpesviridae* was the most strongly enriched for the “stress activated protein kinase signaling cascade” term, which is interesting given that herpesviruses are known to reactivate from latency in response to stress kinase signaling cues;^16^ however, direct host stress kinase phosphorylation of herpesvirus proteins remains poorly studied. Viruses in the *Coronaviridae, Picornaviridae, Adenoviridae, Filoviridae, Togaviridae, Bunyaviridae, Paramyxoviridae, Flaviviridae, Retroviridae,* and *Astroviridae* families were also significantly enriched for stress-kinase motifs, suggesting these viruses have evolved to sense and respond to stress signals during their life cycle. For example, bovine parainfluenza virus in the *Paramyxovirdiae* family is known to co-opt the host p38 MAPK signaling pathway to enhance expression of viral proteins and promote efficient replication.^17^

We also found viruses in the *Retroviridae* and *Hepadnaviridae* families to be enriched for cell cycle-related kinases, a result consistent with the known replication of these viruses in the nucleus. For instance, retroviruses reverse-transcribe their RNA genomes into DNA and integrate them into the host genome as part of their replication cycle, promoting viral persistence in the host. Integration is coordinated alongside the cell cycle stage, occurring primarily in late G1 or early S phase, and several cell cycle-related kinases have been previously implicated in the retrovirus life cycle.^18^ Hepatitis B virus (HBV), a *Hapadnaviridae* member, is known to prefer non-dividing cells arrested in G0/G1 or early S, when the nuclear membrane remains intact, to protect its genome and support replication. In fact, the HBV HBx protein helps maintain cells in G0/G1 or early S phase by modulating host cell cycle regulators.^19^ The extent to which HBV coordinates its life cycle alongside host cell cycle signals remains largely unexplored. Lastly, we found the *Orthomyxoviridae* family, which includes influenza viruses, to be enriched for ion transport-related kinases, which is intriguing given the known reliance of influenza on its viral M2 proton channel to acidify the viral interior during entry and to regulate pH during viral assembly and release.^20^

We next investigated whether virus-kinase interactions were enriched for specific amino acid motif categories relative to humans, since stress and cell cycle motifs are known to be driven by specific motif patterns (i.e., proline at the +1 position)^14^. To assess this, we compared the fraction of high-confidence kinase motifs (KMS > 2) assigned to each motif category, as defined by Johnson et al.^14^, and used the ratio of high-confidence motif occurrence between each virus and human as a metric of selection for specific motif families. Overall, we found viral proteins did not possess overarching selection for particular motif families relative to human proteins (Fig. 2D). However, there was notable variability between viruses, with some viruses showing clear preferences for specific motif classes (Fig. 2D, see outliers). The motif families with greater occurrence in viral relative to human proteins were ULK/TTBK, NEK/ASK, RIPK/WNK, and MAP4K (Fig. 2D). We next evaluated whether specific kinase motif families were preferentially associated with viruses enriched for representative signaling pathways identified in Fig. 2B. For each motif family, we quantified the number of viral peptides from viruses significantly associated with a given pathway relative to all viruses. Pathways related to “regulation of response to stress” and “defense response” showed no motif family preference, whereas the “cell cycle” pathway displayed strong enrichment for divergent CMGC, LKB/CAMKK, and proline-directed kinase motif families (Fig. 2E).

### Viral protein phosphorylation sites are mostly, but not always, surface exposed

We next investigated the relationship between viral protein phosphorylation and the structural context of site occurrence. Specifically, we integrated mass spectrometry phosphoproteomics data from virus-infected cells, 36,643 AlphaFold2-generated viral and human protein structures,^21^ and human kinase motif scores in viral and human proteins (Fig. 3A). We performed global MS phosphoproteomics during Sindbis virus (SINV) infection in HEK293T cells (MOI=1, 24 hours post-infection), KSHV latency reactivation in BC-3G B-cells (20ng/mL phorbol 12-myristate 13-acetate [PMA] for 24 hours), and HIV-1 latency reactivation in JLat 10.6 T-cells (5ug/mL PMA for 24 hours; Fig. S2A and S2B). We detected 43, 91, and 8 high-confidence phosphorylation sites (localization probability>0.75) for SINV, KSHV, and HIV-1, respectively.

**Figure 3.**
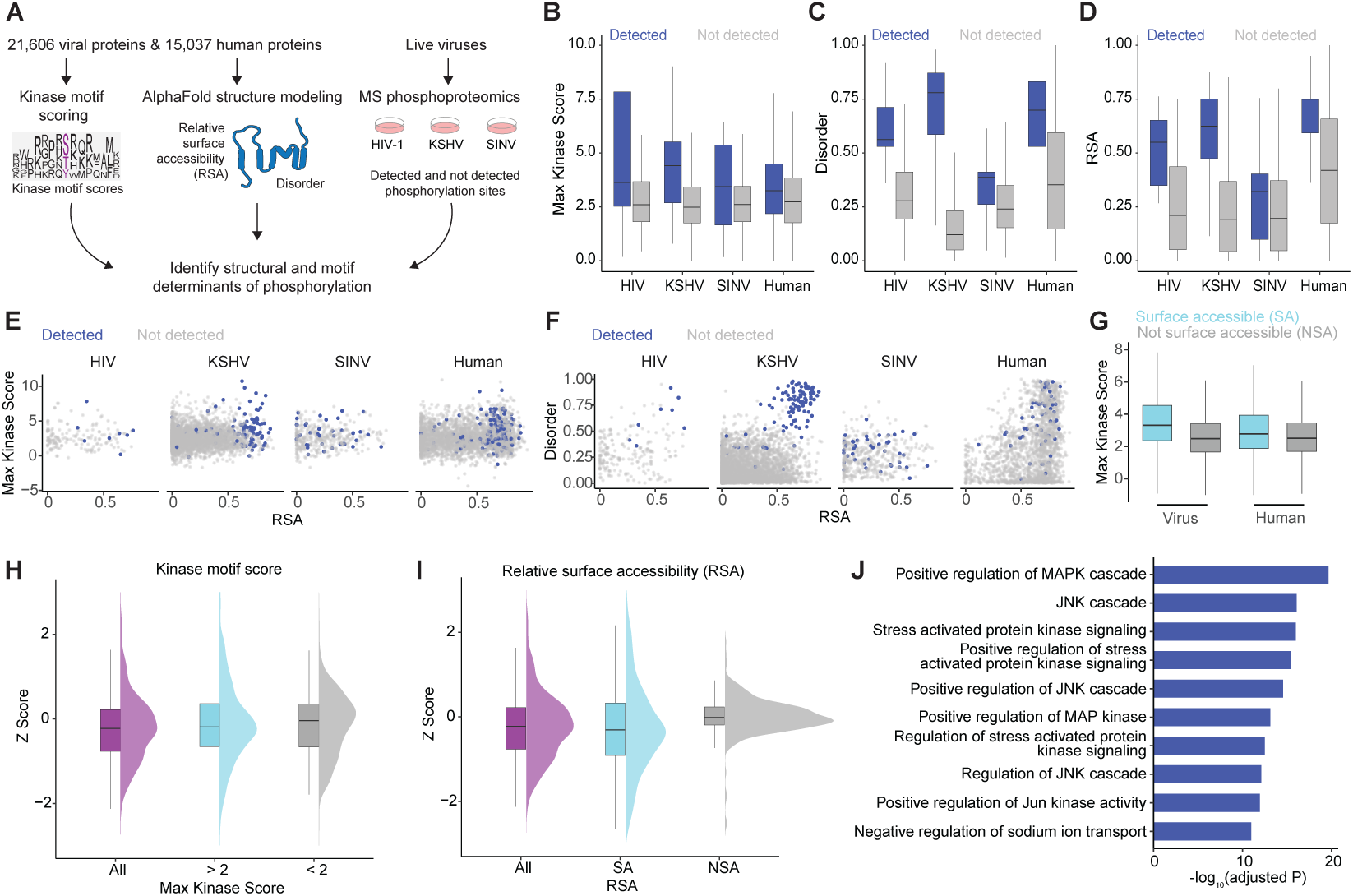
Viral proteins are responsive to phosphorylation largely on their surface. (A) Kinase motifs were annotated for 21,606 viral proteins and 15,037 human proteins. Each protein was modeled with AlphaFold to calculate relative solvent accessibility (RSA) and disorder values. These data were integrated with phosphoproteomics from HIV-1, KSHV, and SINV infections to identify determinants of phosphorylation. (B) Comparison of the maximum kinase motif score per site between phosphoproteomics-detected and undetected sites across HIV, KSHV, SINV, and human proteins. (C) Comparison of intrinsic disorder per site between detected and undetected sites. (D) Comparison of relative surface accessibility (RSA) per site between detected and undetected sites. (E) Relationship between RSA and maximum kinase motif score across all sites and data sources, colored by detection in phosphoproteomics data. (F) Relationship between RSA and intrinsic disorder across sites and data sources, colored by detection in phosphoproteomics data. (G) Comparison of maximum kinase motif scores per site between viral and human proteomes, colored by surface accessibility. Surface accessible (SA) sites are defined as RSA > 0.5; non-surface accessible (NSA) sites are defined as RSA < 0.5. (H) Z-scores calculated per virus–kinase pair, thresholded by whether the maximum kinase motif score was greater or less than 2. (I) Z-scores calculated per virus–kinase pair, thresholded by RSA value above or below 0.25 (SA and NSA, respectively). (J) Top GSOA term enrichment results from Z-scores derived from surface-accessible motifs.

Sites detected during infection exhibited higher maximum kinase motif scores (Fig. 3B) and were more likely to occur in intrinsically disordered regions (Fig. 3C). These sites also tended to be more surface exposed, as indicated by higher relative solvent accessibility (RSA) values (Fig. 3D). However, the magnitude of these trends varied across viruses: while KSHV phosphosites were overwhelmingly disordered and surface accessible, SINV phosphosites showed minimal differences between detected and undetected sites.

We next assessed the relationship between kinase motif scores, disorder, RSA, and phosphorylation. We observed a trend of increasing kinase motif scores associated with surface-exposed phosphorylation sites (Fig. 3E). We also found most detected sites on KSHV and HIV proteins to be on disordered and surface exposed residues (Fig. 3F). Surprisingly, SINV phosphosites were generally more ordered and buried within the protein core, which was also found to be the case for many human sites (Fig. 3F). This may suggest that these sites are phosphorylated during or prior to protein folding. Interestingly, we noticed the proportion of phosphorylated phospho-acceptors was much higher for SINV (10.5%) than for KSHV (3.3%), HIV (4.6%), or human proteins (5.0%), despite all having an equivalent fraction of high-confidence kinase motifs. The high fraction of phosphorylation sites in SINV may reflect their location in non– surface-accessible regions, limiting access by host phosphatases and suggesting a potential role in maintaining viral protein structural integrity.

### Human kinase specificity for viral proteins is enhanced at surface-exposed regions

To further investigate the structural constraints of viral protein phosphorylation by human kinases, we expanded our analysis beyond SINV, KSHV, and HIV-1 to cover 1,391 viruses, comprising 21,606 viral protein structures^21^. We observed phosphorylation sites on the surface (RSA > 0.5) of viral and human proteins exhibited higher max KMS than those located in buried regions (RSA < 0.5; Fig. 3G), suggesting that surface-exposed phosphoacceptors are under stronger selection to evolve high-scoring kinase motif contexts.

We next sought to identify features that determined viral biases toward specific host kinase signaling pathways, relative to the baseline preference of human kinases for human substrates. First, we used the z-score-based analysis outlined above (Fig. 2A) and filtered by max kinase score, but observed no ability to distinguish viral bias towards specific human kinases, as evidenced by equivalent z-score distributions among high- and low-scoring kinase motif thresholds (Fig. 3H; Fig. S2C, top). However, filtering by RSA revealed a bias of viral phosphorylation motifs for certain human kinases. Specifically, the differences between viral and human motif preferences was more pronounced for surface-accessible (RSA>0.25) than non-surface-accessible (RSA<0.25) sites, as indicated by the narrower z-score distribution for kinases predicted for non-surface-accessible sites (Fig. 3I; Fig. S2C, bottom). This suggests that viral protein surfaces experience stronger selection to favor specific kinases. Among the kinases with the highest median z-scores across viruses for surface-exposed sites were RIPK2, IRAK1, and MAP3K7, all known for roles in modulating the stress response (Fig. 3I; Fig. S1B). To globally interrogate this trend, we performed GSOA on kinases with a significant preference for viral proteomes (z>0 & p<0.05), which confirmed enrichment for stress signaling pathways (Fig. 3J; Table S3). Since many stress kinases are known to phosphorylate motifs with a proline in the +1 position, we next systematically evaluated whether amino acid composition at surface-exposed regions could account for the observed kinase preferences. However, we found no significant difference in the surface preference of proline residues between viral and human proteins (Fig. S1C). Similarly, there was no overall enrichement in proline abundance in viral proteomes compared to human proteomes (Fig. S1D). However, we did observe a statistically significant preference for R, Q, D, H, Y, and C on the surface of viral proteins and for L, M, A, G, and S on the surface of human proteins (Fig. S1C).

### Human kinase motifs in viral proteins show positive selection

We next developed a computational pipeline to evaluate whether kinase motifs in viral proteins were under positive selection. Positive selection refers to evolutionary pressure that favors beneficial amino acid changes, typically detected as an excess of nonsynonymous over synonymous substitutions, and can involve repeated toggling between residues that confer an adaptive advantage.^21^ We matched nucleotide sequences to viral proteins in the Nomburg et al.^22^ dataset, yielding 10,629 DNA sequences (Fig. 4A), which we hierarchically clustered based on sequence similarity (see Methods). To identify sequence groups indicative of divergent evolution, we required proteins within a cluster to share at least 70% sequence identity and be a member of the same viral family, requiring at least four proteins per cluster. This resulted in 162 clusters, representing the *Poxviridae* and *Herpesviridae* families (Fig. 4B; Table S4), ranging from four to 12 proteins per cluster (Fig. 4C). As expected, intracluster sequence similarity far exceeded intercluster similarity (Fig. 4D). Codon-based multiple sequence alignment was then performed within each cluster, and aligned sequences were analyzed using the HyPhy suite with the mixed effects model of evolution (MEME) to detect episodic positive selection.^23^

**Figure 4.**
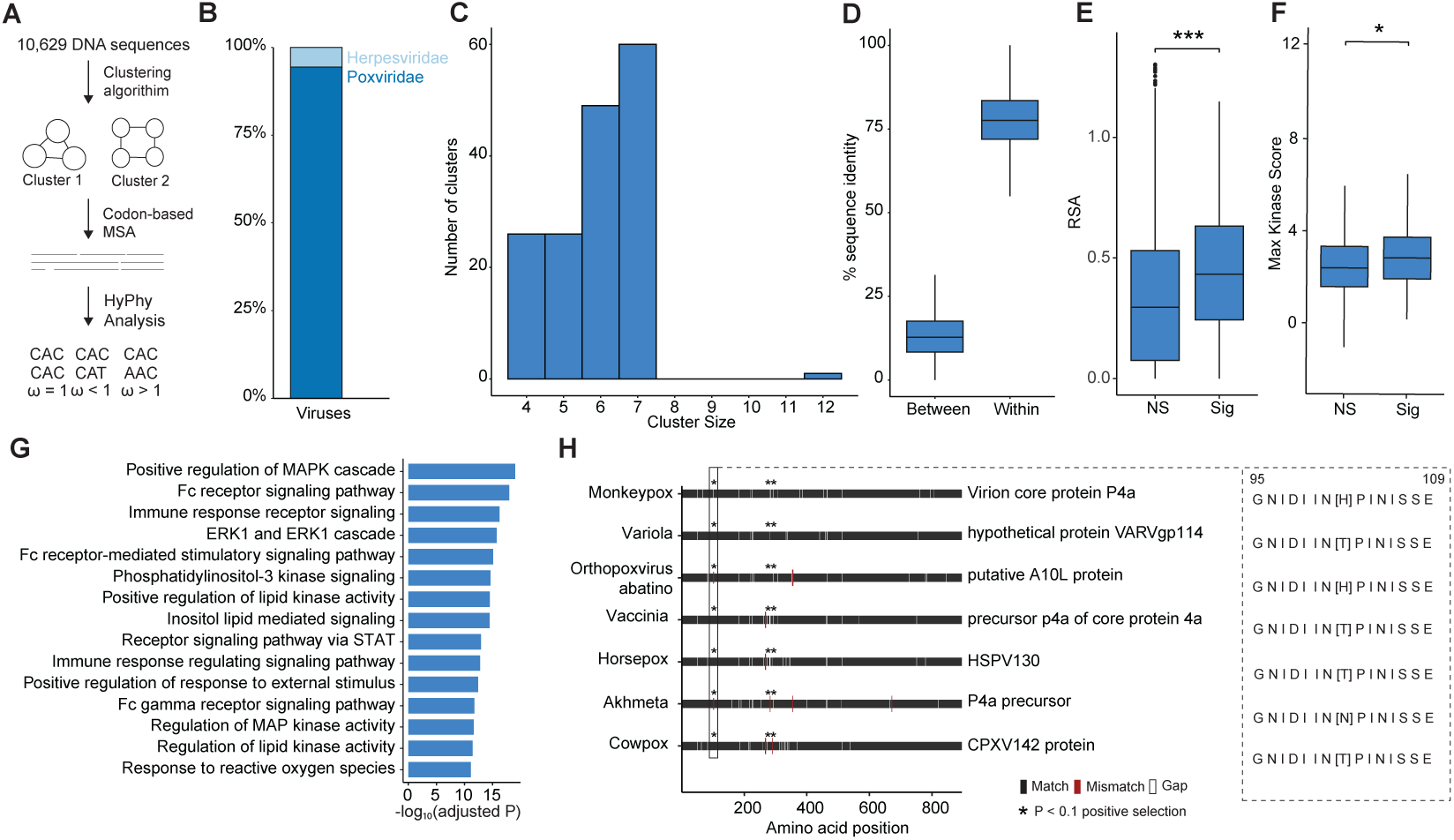
Landscape of motifs under positive selection. (A) 10,692 viral DNA sequences were clustered and subjected to codon-based multiple sequence alignment, followed by HyPhy analysis on individual codons. (B) Distribution of viral families included in the positive selection analysis. (C) Sizes of clusters analyzed for positive selection. (D) Percent sequence similarity of protein sequences within and between clusters. (E) Relative surface accessibility (RSA) of amino acids under significant (Sig) positive selection versus non-significant (NS) amino acids (Wilcoxon rank-sum test, p < 0.001). (F) Maximum kinase scores of phosphoacceptor sites under significant positive selection compared to non-significant sites (Wilcoxon rank-sum test, p < 0.05). (G) Gene Set Overrepresentation Analysis (GSOA) of kinases with elevated scores at sites under significant positive selection. (H) Representative alignment of sequences from one cluster, highlighting a kinase motif across poxviruses.

Prior research found that residues under positive selection were enriched at solvent-exposed sites that influenced protein-protein interactions.^24^ Corroborating prior work, and the validity of our analysis, we also observed residues under positive selection (p < 0.1) exhibited significantly higher solvent-accessibility, reflected by higher RSA scores (Fig. 4E; Fig. S3A; Table S4). We next sought to evaluate whether human kinase motifs on viral proteins were under positive selection. Interestingly, we observed that phosphoacceptors (i.e., S/T/Y) under positive selection had significantly higher maximum kinase motif scores compared to other sites (Fig. 4F). This finding suggests that phosphoacceptor sites under positive selection are enriched for features that elevate kinase motif scores, indicating an evolutionary bias toward retaining the regulatory feedback from host signaling cues. To identify kinase pathways contributing to this enrichment, we identified kinases with higher motif scores on phosphoacceptor residues under positive selection using a z-test approach, followed by GSOA. The resulting pathways were predominantly involved in MAPK cascades and immune signaling (Fig. 4G; Fig. S3B; Table S4), indicating that viral phosphosites are selectively tuned for phosphorylation by host pathways critical to infection.

As an illustrative example, we examined a cluster of seven poxvirus proteins (Fig. 4H) and identified a motif in which the surrounding sequence was fully conserved but the central phosphoacceptor was under positive selection, producing a strong MAPK8 (JNK1) motif (max KMS of 4.57). This example shows that viruses can evolve new phosphoacceptor sites with high-affinity host kinase motifs. Additionally, we examined kinase score distributions for sites where the phosphoacceptor was perfectly conserved within the cluster, but the adjacent motif contained at least one site under significant positive selection. Interestingly, cluster #573 showed a wide kinase score range for MAPK13 (p38-delta) and CDK4-6, with some motifs favoring phosphorylation by MAPK and others by CDK4-6 (Fig. S3C). This suggests that positive selection at the motif level (i.e., change to motif surrounding the phosphoacceptor) can drive divergence in kinase phosphorylation preferences and, potentially, host signaling responsiveness. Taken together, our positive selection analysis reveals that viral proteins preferentially evolve phosphoacceptor sites with strong kinase recognition potential, particularly within MAPK and immune pathways, underscoring phosphorylation as a focal point of viral adaptation.

### Dynamic regulation of Sindbis virus phosphorylation and MAPK signaling reveals a critical dependency

We next sought to probe the impact of human kinase phosphorylation of viral proteins on viral replication. We selected SINV as a model system due to its similarity to more pathogenic alphaviruses such as chikungunya virus.^25^ We first performed MS abundance proteomics and phosphoproteomics of infected HEK293T cells over time (4, 8, 12, 16, 20, and 24 hours post infection) to characterize the dynamics of host kinase signaling activities and viral protein phosphorylation trends over time (Fig. 5A; Table S5). As expected, our data showed a steady increase in viral titer levels by plaque assay (Fig. 5B), viral protein abundance (Fig. 5C) and phosphorylation (Fig. 5D) across the time course. Interestingly, all nine viral proteins were detected in the abundance proteomics data and phosphorylation was detected on all viral proteins except the 6K structural protein (Fig. 5C-D; Fig. S4A-B).

**Figure 5.**
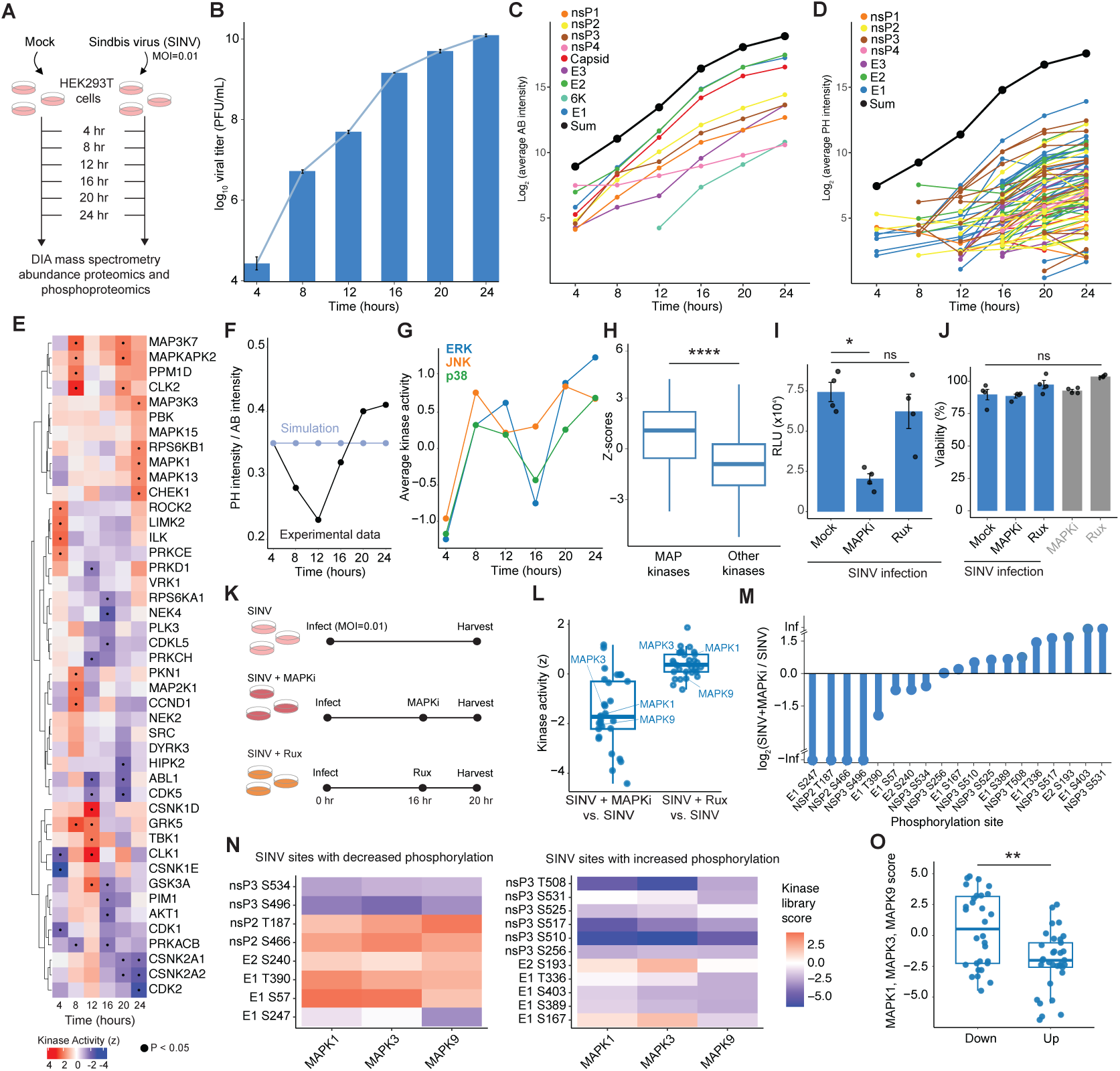
Time-course proteomic profiling and functional validation of host kinase responses during SINV infection. (A) HEK293T cells were infected with SINV (strain TE5’2J/GFP) at an MOI of 0.01, cell lysates were collected at 4 hour time intervals for DIA mass spectrometry abundance and phosphoproteomics. (B) Plaque assay results from SINV time course experiment. (C) Change in viral protein abundance over time, averaged across biological replicates. (D) Change in phosphorylation levels on viral proteins over time, averaged across biological replicates. Each line represents a different phosphopeptide. (E) Changes in host kinase activity over the time course compared to mock-infected controls (F) Ratio of phosphorylation intensity to protein abundance over time (black), compared to model-predicted values. (G) Average kinase activity over time for kinases within the ERK, JNK, and p38 pathways. (H) Z-scores calculated for MAPK kinases in Sindbis virus compared to all other kinases (Wilcoxon rank-sum test, p < 0.0001). (I) Luciferase assay results from SINV-infected cells pre-treated with 1 uM MAPK inhibitors or 1 uM ruxolitinib one hour before infection (Wilcoxon rank-sum test, p < 0.05). Cells were harvested at 24 hours post-infection. (J) Cell viability for each treatment group; 100% represents non-infected, untreated mock controls. (K) HEK293T cells were infected with SINV (strain Toto 1101) at an MOI of 0.01. A MAPK inhibitor cocktail or ruxolitinib was added at 16 hours post-infection, and cells were harvested at 20 hours for proteomic and phosphoproteomic analysis. (L) Kinase activity compared across SINV-infected cells with or without MAPK inhibitor or ruxolitinib treatment; contrasts are shown relative to untreated SINV infection. (M) Detection of phosphorylation sites in MAPK inhibitor-treated versus mock-treated cells, normalized to protein abundance in each condition; values greater than 0 indicate enrichment in MAPK inhibitor-treated cells. (N) Kinase Library scores for sites with increased or decreased phosphorylation, calculated for MAPK1, MAPK3, and MAPK9. (O) Comparison of Kinase Library scores for MAPK1, 3, and 9 across sites with differential phosphorylation (Wilcoxon rank-sum test, p < 0.01).

To assess the impact of infection on host signaling, we estimated host kinase activities over time from the global phosphoproteomics data by using prior knowledge networks of kinase-substrate interactions. This approach allowed us to monitor a consistent set of phosphorylated peptides for each kinase over time (Fig. 5E; Table S6). At 4 hours post infection (hpi), we noted high ROCK2 and LIMK2 activity, both cytoskeleton regulators, possibly reflecting changes that occur during viral entry. Interestingly, at 8 and 12 hours, we noted activation of MAPK and TBK1 stress and innate immune signaling, which was blunted by 16 hpi but resurged by 24 hpi (Fig. 5E).

We next investigated the ratio of viral protein phosphorylation to viral protein abundance to determine whether the levels of viral protein phosphorylation increased linearly or non-linearly with total protein levels. To establish a baseline, we simulated an ODE model of kinase-substrate interactions using first-order mass-action kinetics, which confirmed a constant phosphorylation-to-abundance ratio over time as substrate increases as long as the levels or activities of enzymes do not change (Fig. 5F, blue line; see Methods). In our data, we observed the ratio of phosphorylated to unphosphorylated viral proteins decreased through 12 hours and increased thereafter until 24 hours, suggesting that additional regulatory factors, such as changes in host kinase activities, may be contributing to this trend (Fig. 5F, black line).

To further explore the activity of host kinases throughout infection, we examined the mean activity of MAP kinases over time, grouped by the JNK, ERK, and p38 subfamilies. This analysis confirmed a similarly biphasic activity dynamic, with peaks observed at 8 and 24 hours (Fig. 5G). To extend this analysis, we examined whether sites in the SINV proteome showed stronger preference for MAPK-mediated phosphorylation. Using the z-scores previously generated for a panel of MAPK kinases, we found that these SINV sites had significantly higher z-scores compared to sites associated with other kinases, indicating potential selective pressure for phosphorylation by MAPKs (p < 0.0001; Fig. 5H; Fig. S4C).

We next tested the functional role of MAPK signaling in infection by using a SINV luciferase reporter virus where the luciferase protein is expressed as a fusion with the viral nonstructural protein 3 (nsP3). HEK293T cells were pretreated for one hour with either a DMSO control, 1 μM MAPK inhibitor cocktail (MAPKi: 1 μM doramapimod [p38 inhibitor], 1 μM JNK-IN-8 [JNK inhibitor], and 1 μM U-0126 [MEK inhibitor]), or 1 μM ruxolitinib (Rux: JAK/STAT pathway inhibitor), followed by infection with SINV strain Toto1101/Luc at a multiplicity of infection (MOI) of 1 for 24 hours. We observed that MAPK inhibition significantly reduced SINV genome replication, which was not observed for ruxolitinib-treated cells (Fig. 5I), an effect that could not be explained by differences in cell viability (Fig. 5J).

We next sought to evaluate whether MAPK could directly phosphorylate SINV proteins. To this end, we infected cells with SINV (MOI=0.01, as in Fig. 5A) and added either the MAPK inhibitor cocktail or ruxolitinib at 16 hours post infection for 4 hours (Table S5), at which point cell lysates were collected for abundance proteomics and phosphoproteomics analysis (Fig. 5K; Fig. S4D). As anticipated, kinase-substrate network analysis showed decreased MAPK activity under MAPK inhibitor treatment, with ruxolitinib leaving MAPK activity unchanged (Fig. 5L). Nineteen (19) viral phosphorylation sites were detected in this experiment, eight and eleven of which depicted decreased or increased abundance-normalized phosphorylation, respectively (Fig. 5M). Interestingly, sites showing decreased phosphorylation exhibited higher Kinase Library scores for MAPK1, MAPK3, and MAPK9 of the ERK and JNK signaling pathways (Fig. 5N-O; Fig. S4E), suggesting ERK and JNK kinases directly phosphorylate SINV proteins. Collectively, our results indicate that alphaviruses have evolved to leverage ERK and JNK phosphorylation, and that these interactions are required for efficient viral replication.

## DISCUSSION

Our study revealed a striking enrichment of human kinase motifs across viral proteomes, suggesting that viruses have evolved to biochemically sense host signaling pathways through post-translational modifications. By combining computational sequence-based kinase motif analysis, AI-based modeling of viral protein structures, and deep MS phosphoproteomics profiling of infected cells, we found viral proteins harbor phosphorylated human kinase motifs predominantly recognized by stress, inflammation, and cell cycle kinases. Structural analysis further showed that these phosphorylation sites are often located on protein surfaces, where motif-specificity differences between human and viral proteins are most pronounced. Lastly, inhibiting MAPK signaling during Sindbis virus infection disrupted both viral replication and phosphorylation of phosphorylation sites with high-scoring ERK and JNK kinase motifs. Collectively, our results suggest that phosphorylation is not always a barrier for viruses to overcome, but a potential mechanism by which many viruses sense and respond to specific host cues, allowing them to adapt to the host intracellular environment.

One key class of signals viruses appear to monitor is the cellular stress response. Stress-activated kinases such as p38, JNK, and ERK are frequently activated in infected or damaged cells,^26^ and our analysis identified an abundance of their target motifs within viral proteins. For viruses, sensing stress may provide a critical cue to shift from replication to assembly and egress, especially if the host cell is undergoing apoptosis or immune activation. A dying or distressed cell is a poor environment for completing a replication cycle, but may be optimal for producing and releasing progeny virions before cell death. Along these lines, herpesviruses are known to reactivate from latency in response to p38, ERK, and JNK kinase signaling, producing infectious virions to escape dying cells and infect neighboring healthy cells.^27^ Our findings suggest that many viruses may use these stress kinase signals as a form of temporal regulation to orchestrate their life cycle in synchrony with host cell viability.

In addition to stress, cell cycle kinases, such as cell cycle dependent kinases (CDKs), also possess an abundance of high-confidence motifs in viral proteins. The cell cycle profoundly impacts the biochemical landscape of a cell, including the availability of dNTPs, nucleotide biosynthesis, chromatin accessibility, and replication machinery, all of which are resources that viruses exploit during their replication cycles.^28,29^ We found certain viral families to be enriched for cell cycle kinase motifs, including retroviruses, which are known to exploit the host nucleus for replication.^30^ The lentivirus subfamily of retroviruses integrate their genomes into the host genome, which requires coordination with host cell cycle activities and DNA replication machinery.^31^ Encoding motifs for cell cycle kinases may enable viruses to “read” the cell cycle state and tune their replication cycle to maximize fitness.

There are several limitations to our study. While our motif analysis and phosphoproteomic data suggest widespread kinase engagement across viral proteomes, definitive mechanistic studies will require precise mapping of phosphorylation site functionality on a case-by-case basis. Second, kinase motif prediction algorithms often fail to capture true in vivo substrate specificity, as they consider catalytic site affinity and neglect additional substrate features such as docking domains or scaffold interactions.^32^ Targeted perturbation studies will be needed to definitely link viral protein phosphorylation events to their upstream kinases in vivo.

Importantly, our findings open several translational avenues. For example, therapeutics could deliberately reprogram host signaling to disrupt viral life cycle transitions, reducing replication and allowing time for mounting an immune response. Moreover, understanding how individual patient signaling responses affect viral life cycle progression could help to explain variability in disease severity and outcomes. Ultimately, this work lays a systems-level foundation for how viruses exploit dynamic cellular states to guide their life cycle, reframing viruses not as indiscriminate replicators but as entities that actively sense and respond to their intracellular environment.

## METHODS

### COMPUTATIONAL METHODS

#### Curation of viral proteomes from Expasy

Viral protein sequences for 125 human-infecting viruses were collected from those annotated as human-infecting in Expasy (viralzone.expasy.org), where proteome files were linked to uniprot.org and downloaded as FASTA files. These sequences were then used for downstream kinase motif analysis using the Kinase Library.

#### Curation of viral proteomes from Nomburg et al

Structural data in the form of PDB files for 21,606 viral proteins from 1,391 viruses were collected from publicly available Zenodo records.^22^ The PDB files were downloaded and stored locally. An R script was used to extract the RefSeq Protein ID from the file names, where it appears as an underscore-delimited string.

#### Kinase motif analysis

Fifteen-mer sequences centered on all serine, threonine, and tyrosine residues were generated from the human proteome, 125 human-infecting viral proteomes, and the 21,606 viral proteomes extracted from Nomburg et al.^22^ Each phosphoacceptor site was scored against the human Kinase Library, using serine/threonine kinases for S/T sites and tyrosine kinases for Y sites. Scores reflect predicted kinase and substrate compatibility, where 0 represents a random association and positive or negative values indicate favorable or unfavorable interactions. This analysis produced a table that assigns a kinase motif score (KMS) to each phosphosite and kinase pair.

#### Calculation of relative surface accessibility (RSA) metric

Accessible surface area (ASA) for each residue was calculated using the Define Secondary Structure of Proteins (DSSP) tool on PDB files curated from Nomburg et al.^22^ and on structures generated independently with AlphaFold2 for HIV-1, KSHV, and SINV proteins. ASA values were then normalized by the maximum solvent-accessible surface area for each amino acid type, as reported by Tien et al.^32^ to calculate the relative surface accessibility (RSA) for each residue.

#### Defining high-confidence virus-kinase interactions

For each kinase, we compared viral kinase motif scores to human kinase motif scores for the same kinase across both the human-infecting viral proteome dataset and the dataset from Nomburg et al.^22^ Although the human proteome had fewer proteins, they had more phosphoacceptors. To control for differences in sample size, human scores were randomly downsampled to match the number of viral scores for each kinase. Analyses were only performed when at least 30 phosphoacceptors were available for both groups, in accordance with the Central Limit Theorem. A z-score and p-value were then calculated for each virus-kinase pair, representing the difference between viral and human kinase motif scores. Significant virus-kinase interactions were defined as those with a z-score greater than zero and a p-value less than 0.05. In addition, we required that each virus contain at least seven high-confidence sites for the kinase, where high confidence was defined as a motif score greater than two. This filtering process yielded a table of significant virus-kinase pairs for downstream analysis.

#### Gene set overrepresentation analysis of virus-associated kinases

Gene set overrepresentation analysis (GSOA) was performed using the curated list of significant virus-kinase interactions. For the human-infecting viral proteome dataset, we selected the top 50 kinases for each virus (or all available if less than 50), ranked based on z-score, and performed GSOA using the Gene Ontology Biological Processes database acquired from MSigDB. The background universe was defined as the HGNC names of all kinases included within the Kinase Library algorithm.^14^ Even though we controlled for kinases as the background, we still found enrichment for genetic phosphorylation terms. The following GO biological process terms were excluded from the analysis to avoid redundancy among kinase-related categories: *protein autophosphorylation*, *positive regulation of protein serine/threonine kinase activity*, *positive regulation of protein kinase activity*, *positive regulation of kinase activity*, *regulation of protein serine/threonine kinase activity*, *peptidyl-serine modification*, *peptidyl-threonine modification*, *activation of protein kinase activity*, and *peptidyl-tyrosine modification*. which we manually removed from the visualization. Pathway terms were ranked by the number of viruses associated with kinases showing a statistically significant enrichment with that term at an adjusted p-value less than 0.05. We also performed this analysis at the viral family level by selecting the top 50 kinases for each family (or all available if less than 50), also ranked by Z-score, and applying the same GSOA approach. For the viral proteome dataset from Nomburg et al.^22^ (Fig. 3), we used z-scores calculated for sites with RSA values greater than 0.25 and ranked kinases by their median z-score across viruses within each family. The top 50 kinases were selected for GSOA.

#### Categorization of kinase motif classes in viral and human proteins

For each virus in the Expasy dataset, we calculated the proportion of high-confidence motifs (KMS > 2) within each motif class as defined by Johnson et al.¹⁴ The same analysis was performed for the human proteome. We then computed the log₂ ratio of the proportion of high-confidence motifs in each virus relative to human (Fig. 2D). To assess whether viruses enriched in specific signaling pathways also exhibited enrichment in particular motif classes, we focused on those significant for “regulation of response to stress,” “defense response,” and “cell cycle.” For these viruses, the proportion of high-confidence motifs was recalculated as described above and normalized to the proportion observed across all viruses in the dataset (Fig. 2E).

#### AlphaFold2 analysis for human proteins, HIV-1, KSHV, and SINV

Protein structures for the human, HIV-1, KSHV, and SINV proteomes were generated using ColabFold v1.5.5 with MMseqs2 Batch mode. FASTA sequences were input into ColabFold with five models and three recycles per prediction. The resulting PDB files were processed using DSSP to calculate accessible surface area (ASA) values for each residue, which were then used to compute relative surface accessibility (RSA).

### Calculation of disorder metric

Intrinsically disordered regions were identified by calculating disorder scores for each residue using the IUPred2A algorithm, applied to a subset of proteins from the human proteome and the viral proteomes from Nomburg et al.^22^ These disorder scores were integrated with the relative surface accessibility (RSA) and kinase motif score metrics for downstream analysis.

#### Integration of kinase motif analysis and mass spectrometry phosphoproteomics data

We performed in silico digestion of the HIV-1, KSHV, SINV, and human proteomes using the cleaver package version 1.38.0, simulating trypsin digestion. Peptides were filtered to include fragments between seven and 52 amino acids in length. Whenever comparing our kinase motif analyses to phosphoproteomics datasets, we used only phosphoacceptors that occurred on tryptic peptides.

#### Mass spectrometry proteomics data search and quantitative analysis

Mass spectra from each DIA dataset were searched against a database consisting of Uniprot Homo sapiens sequences downloaded on October 6, 2023, along with HIV-1 (HIV/R7/E^-^/GFP)^34^, KSHV, and SINV (Toto 1101) proteomes. For protein abundance samples, data were searched using the default Biognosys settings, with methionine oxidation as a variable modification, carbamidomethylation of cysteine as a static modification, and a final false discovery rate of 0.01 applied at the peptide, peptide spectrum match, and protein levels. For phosphopeptide-enriched samples, Biognosys settings were modified to include phosphorylation of serine, threonine, and tyrosine as variable modifications.

Quantitative analyses were performed in R. Initial quality control, including inter-run clustering, correlation analyses, principal component analysis (PCA), and assessment of peptide and protein counts and intensities, were conducted using the MSstats package version 4.16.1. Statistical analysis of phosphorylation and protein abundance changes between mock- and virus-infected samples was performed using peptide ion fragment data from Spectronaut processed through MSstats. Phosphorylation site quantification was conducted using MSstats, applying site conversion and quantification with default settings. Peptides containing the same set of phosphorylated sites were grouped and quantified as phosphorylation site groups.

For both phosphopeptide and protein abundance pipelines, MSstats performed normalization by median equalization without imputation of missing values. Summarization was performed by median smoothing (Tukey’s median polish) to combine intensities from multiple peptide ions or fragments into a single intensity value for each protein or phosphorylation site group. Statistical testing for differences in intensity between infected and control samples was performed using the default MSstats settings; unless otherwise specified, adjusted p-values were calculated using the Student’s t-test with the Benjamini-Hochberg method for false discovery rate (FDR) correction, including cases where sample sizes were limited to two replicates.

#### Positive selection evolutionary analysis

A curated database of 10,629 viral nucleotide sequences was used as input, selected from proteins within Nomberg et al.^22^ Pairwise sequence similarity was calculated with a k-mer-based approach, in which sequences were parsed into overlapping 6-mers. For each pair of sequences, similarity was quantified using the Jaccard index on the sets of unique 6-mers. Pairs with similarity scores ≥70% were retained, and only those belonging to the same viral family were considered for downstream analysis. Retained pairs were represented as weighted edges, with edge weights equal to the percent similarity. To limit network density, the 12 strongest edges per protein were preserved.

Edge weights were then re-scaled using shared-nearest-neighbor overlap, with a minimum boost factor of 0.1. Community detection was performed using the Leiden algorithm at four resolution parameters (0.3, 0.6, 0.9, and 1.2). The resolution yielding the highest number of clusters with ≥4 proteins was selected. Proteins without any retained edges were assigned to singleton clusters. Clusters containing fewer than four sequences were excluded from further analysis.

Nucleotide sequences were translated into amino acid sequences, and proteins within each cluster were aligned using MAFFT. Multiple sequence alignments were then reverse-mapped onto codon sequences to generate codon-level FASTA files for each cluster. Each codon alignment was analyzed with HyPhy MEME^23^ (Linux deployment) to identify sites under episodic positive selection. HyPhy produced cluster-specific JSON output files, which were aggregated into a combined results table. This table was subsequently integrated with previous RSA and kinase motif scoring datasets to generate a master dataset for downstream analyses.

#### Mathematical computational modeling

To mathematically model the conversion of substrate (unmodified viral protein) to product (phosphorylated viral protein) during infection, we utilized a simulated mathematical model (Fig. 5). We modeled the rate of change of the substrate as 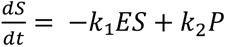 and the rate of change of the product as 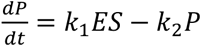. We simulated the mathematical model using the deSolve^35^ package in R to steady state. Values for *S* were set at the total SINV protein abundance at each time point. Constants *k*_1_and *k*_2_ were set to an arbitrary value of 10, while the value for *E* was set to 0.35 so as to be equivalent to the initial experimentally determined ratio.

#### Kinase activity analysis from phosphoproteomics data

We performed kinase activity analysis on phosphoproteomics data collected during HIV-1, KSHV, and SINV infection, as well as during a time course of SINV infection. Log2 fold changes (Log2FC) in phosphosite intensities between infected and mock-treated cells were calculated and used to infer kinase activity states using prior knowledge networks of kinase-substrate interactions derived from the Omnipath database.^36^ Kinase activities were inferred as a z-score from a Z-test comparing Log2FCs in phosphosite measurements of the known substrates (per kinase) against the overall distribution of detected Log2FCs across the sample. This statistical approach has been previously shown to perform well at estimating kinase activities.^37,38,39^

### EXPERIMENTAL METHODS

#### HIV reactivation experiment

J-Lat 10.6 cells were maintained in RPMI-1640 medium (Gibco) supplemented with 10% fetal bovine serum (FBS, Gibco) and 1% penicillin/streptomycin (Gibco) at 37 °C and 5% CO₂. For HIV-1 reactivation, 10 × 10⁶ cells were treated in a 10 cm dish with 5 ng/mL phorbol 12-myristate 13-acetate (PMA) for 24 h, which resulted in 70-90% reactivated cells assayed by GFP expression via flow cytometry.

#### KSHV reactivation experiment

BC-3-G cells were maintained in RPMI-1640 medium (Gibco) supplemented with 15% fetal bovine serum (FBS, Gibco) and 1.5 ug/mL puromycin at 37 °C and 5% CO₂. They were simulated with 20 ng/mL phorbol 12-myristate 13-acetate (PMA) for 24 h, which resulted in ∼40% reactivated cells assayed by GFP expression via flow cytometry.

#### SINV infection experiments

For all Sindbis virus (SINV) infection experiments, 3 × 10⁶ HEK293T cells were seeded per 10 cm dish one day prior to infection. Cells were either mock-infected or infected with EGFP-labeled SINV strain TE5’2J/GFP. Infections were performed in 2 mL of DMEM containing 1% FBS at 37°C for 1 hr to allow virus adsorption. Following adsorption, 8 mL of complete growth medium (DMEM supplemented with 10% FBS and 1% penicillin-streptomycin) was added to each plate to mark the 0 hr time point. For the single time-point SINV phosphorylation experiment, cells were infected at a multiplicity of infection (MOI) of 1 and harvested at 24 hpi. Cells were scraped into the culture medium, collected by centrifugation at 400 × g for 4 min, washed once with 1 mL of ice-cold PBS, and centrifuged again under the same conditions. Cell pellets were lysed in 4% sodium deoxycholate (SDC) lysis buffer, boiled at 95°C for 5 min, and stored at −80°C until further processing. For the SINV time-course, cells were infected at an MOI of 0.01 and collected every 4hr for 24hr post-infection. Following harvest as described above, cell pellets were lysed in 8 M urea buffer and stored at −80°C. For the SINV kinase inhibitor experiment, cells were infected at an MOI of 0.01 and treated at 16 hpi with either ruxolitinib (1 μM, MedChemExpress) or a MAPK inhibitor cocktail consisting of doramapimod (1 μM, MedChemExpress), JNK-IN-8 (1 μM, Selleckchem), and U-0126 (1 μM, Selleckchem). Cells were collected at 20hpi, washed, and lysed in 8M urea buffer as described above before storage at −80°C.

#### Mass spectrometry proteomics sample processing

Generally, unless specified differently below, cells were lysed under denaturing conditions, proteins were reduced and alkylated, and digested for global and phosphoproteomic analysis (experimental specifics to follow). Protein concentrations were quantified by Bradford assay. Disulfide bonds were reduced with 10 mM tris(2-carboxyethyl)phosphine (TCEP) for 30 min and alkylated with 40 mM 2-chloroacetamide (2-CAM) for 30 min at room temperature in the dark. Proteins were digested overnight (16-18 h) with sequencing-grade trypsin (Promega) and, where indicated, Lysyl Endopeptidase (Lys-C, Wako). Post-digestion handling, peptide cleanup, and desalting followed the experiment-specific protocols below. Phosphorylated peptides were enriched using Ti-IMAC HP magnetic beads (Resyn Biosciences) at a peptide-to-bead ratio of 1:2 according to the manufacturer’s protocol. Eluted phosphopeptides were immediately acidified to pH ≈ 3 with 10% formic acid, dried by vacuum centrifugation, and stored at −80 °C until LC– MS/MS analysis.

For SINV timecourse and MAPKi experiments, cells were lysed in 8 M urea, 0.1 M Tris-HCl, 250 mM NaCl, and 50 mM ammonium bicarbonate supplemented with protease (mini-cOmplete, Roche) and phosphatase inhibitors (PhosSTOP, Roche). Lysates were immediately frozen at −80 °C. Upon thawing, DNA was sheared via probe sonication on ice (15 s × 2 cycles, 15% amplitude). Approximately 1 mg of total protein per sample was processed using the general workflow described above. Urea concentration was diluted from 8 M to 2 M with 0.1 M Tris-HCl prior to digestion to permit protease activity. Digestion was performed at 25 °C for 18 h with shaking at 800 rpm using trypsin (1:100, w/w) and Lys-C (1:50) as specified. Following digestion, samples were acidified with 10% TFA to pH 2–3 and clarified by centrifugation (18,000 × g, 10 min, 4 °C). Peptides were desalted using Oasis HLB cartridges (Waters) on a vacuum manifold. Cartridges were activated with 1 mL 80% acetonitrile (ACN)/0.1% TFA, equilibrated three times with 1 mL 0.1% TFA, and samples were loaded twice. Columns were washed three times with 1 mL 0.1% TFA and peptides were eluted with 800 µL 50% ACN/0.25% formic acid. Approximately 10% of each sample was retained for global proteomics; the remainder underwent Ti-IMAC enrichment as described in the general workflow.

For the HIV experiment, cells were pelleted at 400 × g for 5 min at 4 °C, washed twice with ice-cold 1× PBS (Corning), and lysed in 300 µL of 4% sodium deoxycholate, 0.1 M Tris-HCl (pH 8). Lysates were sonicated on ice (15 s × 2 cycles at 10–15% amplitude), boiled for 5 min at 95 °C, and clarified by centrifugation (18,000 × g, 10 min, 4 °C). Protein aliquots corresponding to 1 mg were normalized to 200 µL and processed according to the general workflow. Protein cleanup and on-bead digestion were performed using Sera-Mag SpeedBeads (Cytiva; 1:1 hydrophilic/hydrophobic mixture) at a 1:10 protein-to-bead (w/w) ratio in 50% ethanol (v/v) for 15 min at 23 °C (1200 rpm). After three washes with 80% ethanol, beads were resuspended in 0.1 M ammonium bicarbonate (pH 8) and digested overnight (16–18 h, 37 °C, 1200 rpm) with trypsin (1:100) and Lys-C at a 1:100 (w/w) enzyme-to-substrate ratio. Following digestion, peptides were collected, brought to 0.1% TFA, clarified by centrifugation, and dried. Approximately 10% of each sample was retained for global proteomics; the remainder underwent Ti-IMAC enrichment as described in the general workflow.

For the KSHV experiment, cells were lysed in 6M guanidine hydrochloride (Sigma Aldrich), boiled at 95°C for 5 minutes, and stored on ice until sonication. Lysates were sonicated on ice (15 s × 2 cycles at 10–15% amplitude), boiled for 5 min at 95 °C, and clarified by centrifugation (18,000 × g, 10 min, 4 °C). Protein aliquots corresponding to 1 mg were normalized to 200 µL and processed according to the general workflow. Digestion was performed using 1:100 trypsin. Following digestion, samples were acidified with 10% TFA to pH 2–3 and clarified by centrifugation (18,000 × g, 10 min, 4 °C). Peptides were desalted using Oasis HLB cartridges (Waters) on a vacuum manifold. Cartridges were activated with 1 mL 80% acetonitrile (ACN)/0.1% TFA, equilibrated three times with 1 mL 0.1% TFA, and samples were loaded twice. Columns were washed three times with 1 mL 0.1% TFA and peptides were eluted with 800 µL 50% ACN/0.25% formic acid. Approximately 10% of each sample was retained for global proteomic analysis, and the remainder was subjected to High Select Fe-NTA phosphopeptide enrichment (Thermo) according to the manufacturer’s instructions.

For the single-time point SINV experiment, cells were lysed in 300 µL of 4% sodium deoxycholate, 0.1 M Tris-HCl (pH 8). Lysates were sonicated on ice (15 s × 2 cycles at 10–15% amplitude), boiled for 5 min at 95 °C, and clarified by centrifugation (18,000 × g, 10 min, 4 °C). Protein aliquots corresponding to 1 mg were normalized to 200 µL and processed according to the general workflow. Digestion was performed using 1:100 LysC and 1:100 trypsin. Following digestion, samples were acidified with 10% TFA to pH 2–3 and clarified by centrifugation (18,000 × g, 10 min, 4 °C). Peptides were desalted using Oasis HLB cartridges (Waters) on a vacuum manifold. Cartridges were activated with 1 mL 80% acetonitrile (ACN)/0.1% TFA, equilibrated three times with 1 mL 0.1% TFA, and samples were loaded twice. Columns were washed three times with 1 mL 0.1% TFA and peptides were eluted with 800 µL 50% ACN/0.25% formic acid. Approximately 10% of each sample was retained for global proteomics; the remainder underwent Ti-IMAC enrichment as described in the general workflow.

For phosphopeptide enrichment, Ti-IMAC HP beads (Resyn Biosciences) were used at a peptide:bead ratio of 1:2 (w/w). Beads were equilibrated three times with binding buffer (0.1 M glycolic acid in 80% acetonitrile, 5% TFA). Dried peptide samples were resuspended in 200 µL binding buffer, added to equilibrated beads, and incubated for 30 min at 23 °C, 1200 rpm. The unbound fraction was discarded. Beads were washed sequentially with 200 µL each of: binding buffer, wash buffer 1 (60% ACN, 1% TFA, 200 mM NaCl), wash buffer 2 (60% ACN, 1% TFA), and LC-MS grade water. Phosphopeptides were eluted twice by incubating beads with 150 µL of 1% (v/v) ammonium hydroxide (in LC-MS grade water) for 10 min at 23 °C, 1200 rpm. Eluates were transferred to a new protein LoBind tube containing 50 µL of 10% (v/v) formic acid, pooled, and dried in a SpeedVac (Labconco). Dried peptides were resuspended in 0.1% formic acid, and approximately 500 ng of protein content was analyzed on a timsTOF HT (Bruker Daltonics).

#### Mass spectrometry proteomics acquisition

Dried peptides were resuspended in 0.1% (v/v) formic acid (FA) in MS-grade water and analyzed on a timsTOF HT mass spectrometer (Bruker Daltonics, positive ion mode) coupled to a Vanquish Neo UHPLC system. Mobile phase A consisted of 0.1% (v/v) FA in MS-grade water and mobile phase B of 0.1% (v/v) FA in 100% MS-grade acetonitrile. The LC was operated in trap-and-elute mode, with peptides first trapped on a PepMap Neo Trap column (5 mm, 100 Å, 5 µm) and separated on an Aurora Elite C18 column (15 cm, 100 Å, 1.5 µm; IonOpticks) maintained at 50 °C using a Bruker CaptiveSpray source (Sonation Lab Solutions) and ionized at 1700 V.

For KSHV phosphoproteomics, peptides were separated at 1 µL/min using a gradient from 5– 35% B over 40 min, then to 45% B over 10 min, and to 95% B over 7 min. For HIV phosphoproteomics, the gradient was 3–16% B over 32 min at 0.3 µL/min, then to 30% B over 20 min, to 45% B over 4 min, to 60% B over 1 min, and to 95% B over 3 min. For SINV phosphoproteomics, peptides were eluted with 3–25% B over 76 min at 0.3 µL/min, then to 45% B over 10 min, to 60% B over 1 min, and to 95% B over 3 min. For SINV time-course abundance samples, gradients were 5–35% B over 37 min at 0.3 µL/min, then to 45% B over 4 min, to 60% B over 1 min, and to 95% B over 3 min. For SINV time-course phosphoproteomics, gradients were 3–16% B over 26 min, then to 30% B over 11.5 min, to 45% B over 4 min, to 60% B over 1 min, and to 95% B over 2.5 min. For SINV inhibitor abundance analyses, gradients were identical to those of the time-course abundance runs (5–35% B over 37 min, etc.), while SINV inhibitor phosphoproteomics used the same gradient as the time-course phosphoproteome (3–16% B over 26 min, etc.).

All samples were analyzed in DIA-PASEF mode with MS1 scans acquired from 100–1700 m/z and a dual-TIMS analyzer ramp rate of 9.42 Hz, with 100 ms accumulation and ramp times (100% duty cycle). For KSHV phosphoproteomics, the ion mobility range was 0.63–1.52 V·s/cm², isolation windows were 40 Da (1 Da overlap) with 25 mass steps per 1.06 s cycle, and collision energy decreased linearly from 59 eV at 1/K₀ = 1.6 V·s/cm² to 20 eV at 1/K₀ = 0.6 V·s/cm², collecting MS/MS spectra from 251.3–1227.3 m/z. For HIV phosphoproteomics, the ion mobility range was 0.68–1.4 V·s/cm² with 28 Da isolation windows (1 Da overlap), 39 mass steps per 1.91 s cycle, and collision energy decreasing from 59 eV at 1.4 V·s/cm² to 20 eV at 0.68 V·s/cm² (MS/MS 297.8–1351.8 m/z). For SINV phosphoproteomics, the range was 0.71–1.41 V·s/cm² with 25 Da isolation windows, 40 steps per 1.59 s cycle, and collision energy decreasing from 59 eV at 1.41 V·s/cm² to 20 eV at 0.71 V·s/cm² (MS/MS 340.9–1301.9 m/z). For SINV time-course abundance, parameters included a 0.66–1.41 V·s/cm² range, 22 Da windows, 49 steps per 1.38 s cycle, and collision energy decreasing from 59 eV at 1.35 V·s/cm² to 20 eV at 0.65 V·s/cm² (MS/MS 267.7–1297.7 m/z). For SINV time-course phosphoproteomics, the range was 0.73–1.45 V·s/cm² with 25 Da windows, 47 steps per 1.59 s cycle, and collision energy decreasing from 59 eV at 1.41 V·s/cm² to 20 eV at 0.71 V·s/cm² (MS/MS 350.0–1479.0 m/z). For SINV inhibitor abundance, settings were 0.67–1.44 V·s/cm² with 24 Da windows, 49 steps per 1.38 s cycle, and collision energy decreasing from 59 eV at 1.41 V·s/cm² to 20 eV at 0.71 V·s/cm² (MS/MS 270.3– 1398.3 m/z). Finally, for SINV inhibitor phosphoproteomics, the ion mobility range was 0.64–1.4 V·s/cm² with 25 Da windows, 47 steps per 1.38 s cycle, and collision energy decreasing from 59 eV at 1.41 V·s/cm² to 20 eV at 0.71 V·s/cm² (MS/MS 273.0–1402.0 m/z).

#### SINV luciferase experiment

A total of 1 x 10^6^ HEK293T cells were seeded per well in 24-well plates. The following day, the culture medium was aspirated, and the cells were treated with either MAPi or ruxolitinib (1 uM) in 150 uL 1% FBS in PBS. After 1 hr incubation with the inhibitors at 37C, SINV (add strain info, the luciferase one), was added directly to the inoculum at an MOI of 1. Following a 1hr virus adsorption period at 37C, 350uL of complete growth medium was added to each well to mark the 0-hr time point. After 24 hr infection, luminescence was recorded using a plate reader following the manufacturer’s protocol for the Promega Luciferase Assay System (Promega).

#### MTT assay during SINV infection experiment

At the indicated time point, the culture medium was replaced with 500 uL of fresh complete medium, and 10 uL of 12mM MTT stock solution (prepared in PBS) was added to each well. Plates were incubated for 2hr at 37 C. 500 uL of SDS-HCl (10% SDS in 0.01M HCl), was added to each well and the plates were incubated for another 4hr. Wells were mixed by pipetting, and the absorbance was measured at 570 nm using a microplate reader. To calculate cell viability, background absorbance from wells containing media and reagents with no cells was subtracted from all measurements. Absorbance values for each condition were then normalized to the average absorbance of untreated control wells to calculate the percentage of viable cells.

#### Quantification of infectious SINV particle production via plaque assay

To determine viral titers, BHK-21 cells were infected at 10-fold dilutions in 1% FBS in PBS and incubated at 37C for 1 hour with rocking of the plate every 15 minutes. An overlay consisting of 2X MEM and 4.5% Avicel was added to each well and the plate was incubated at 37C overnight. 24hr post-infection, cells were fixed with 7% formaldehyde for 30 minutes and stained with 1X crystal violet for 30 minutes. Plates were washed with water and plaques were counted by hand.

## SUPPLEMENTAL FIGURES

**Supplemental Figure 1.**
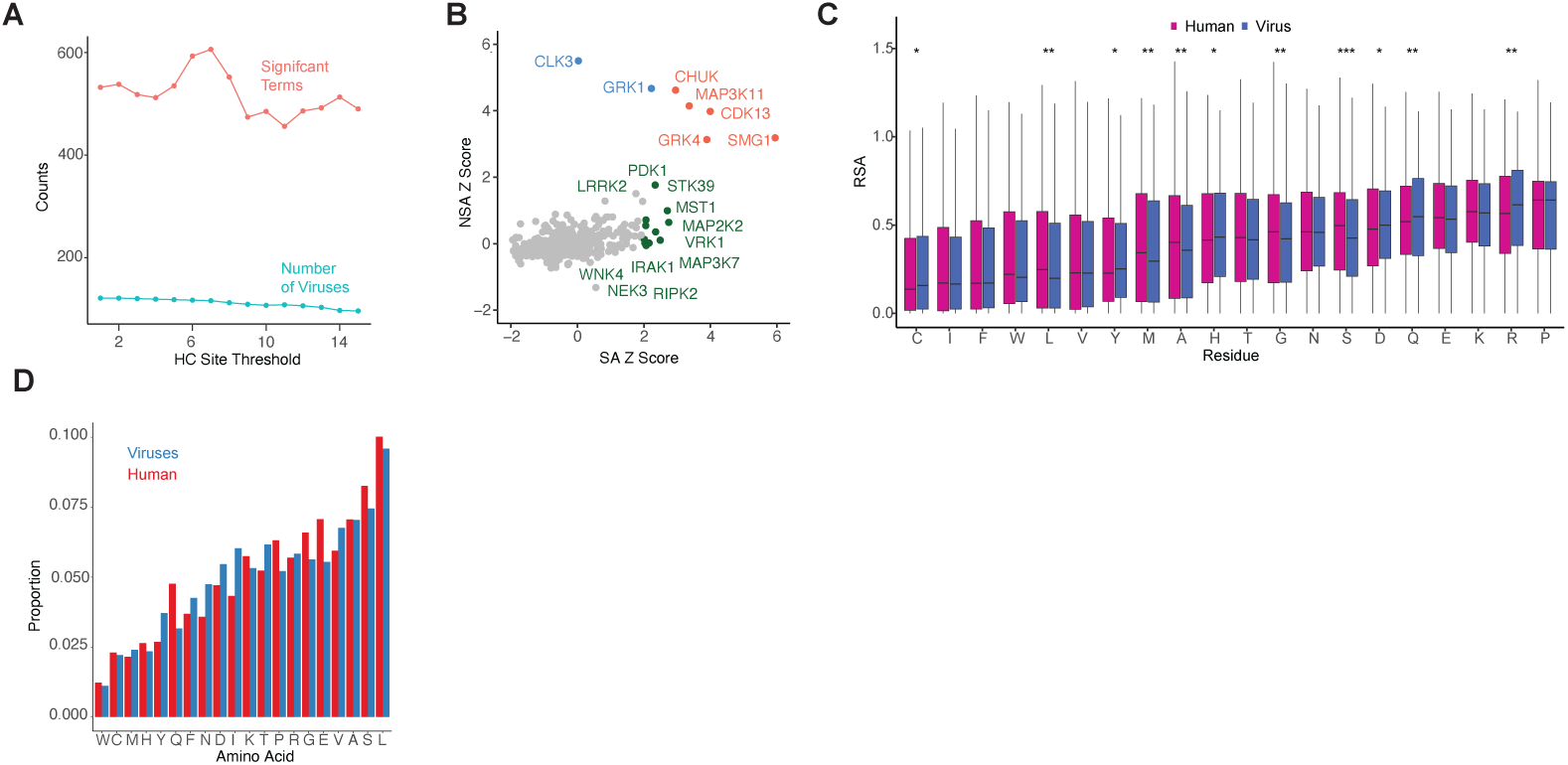
Identification of significant virus-kinase pairs and proteome composition of viruses and humans. (A) Performance of different threshold values for defining high-confidence sites per virus-kinase pair. Red indicates the number of significantly enriched GSOA terms (p < 0.05), while blue shows the number of viruses retained in the GSOA analysis after applying each threshold. (B) Comparison of median Z-scores per kinase when calculated for surface-accessible versus non-surface-accessible sites. Green indicates kinases enriched only in SA sites, blue indicates those enriched only in NSA sites, and orange indicates enrichment in both. (C) Comparison of RSA distributions across viral and human proteomes, stratified by amino acid residue type. (D) Baseline amino acid composition across viral (blue) and human (red) proteomes.

**Supplemental Figure 2.**
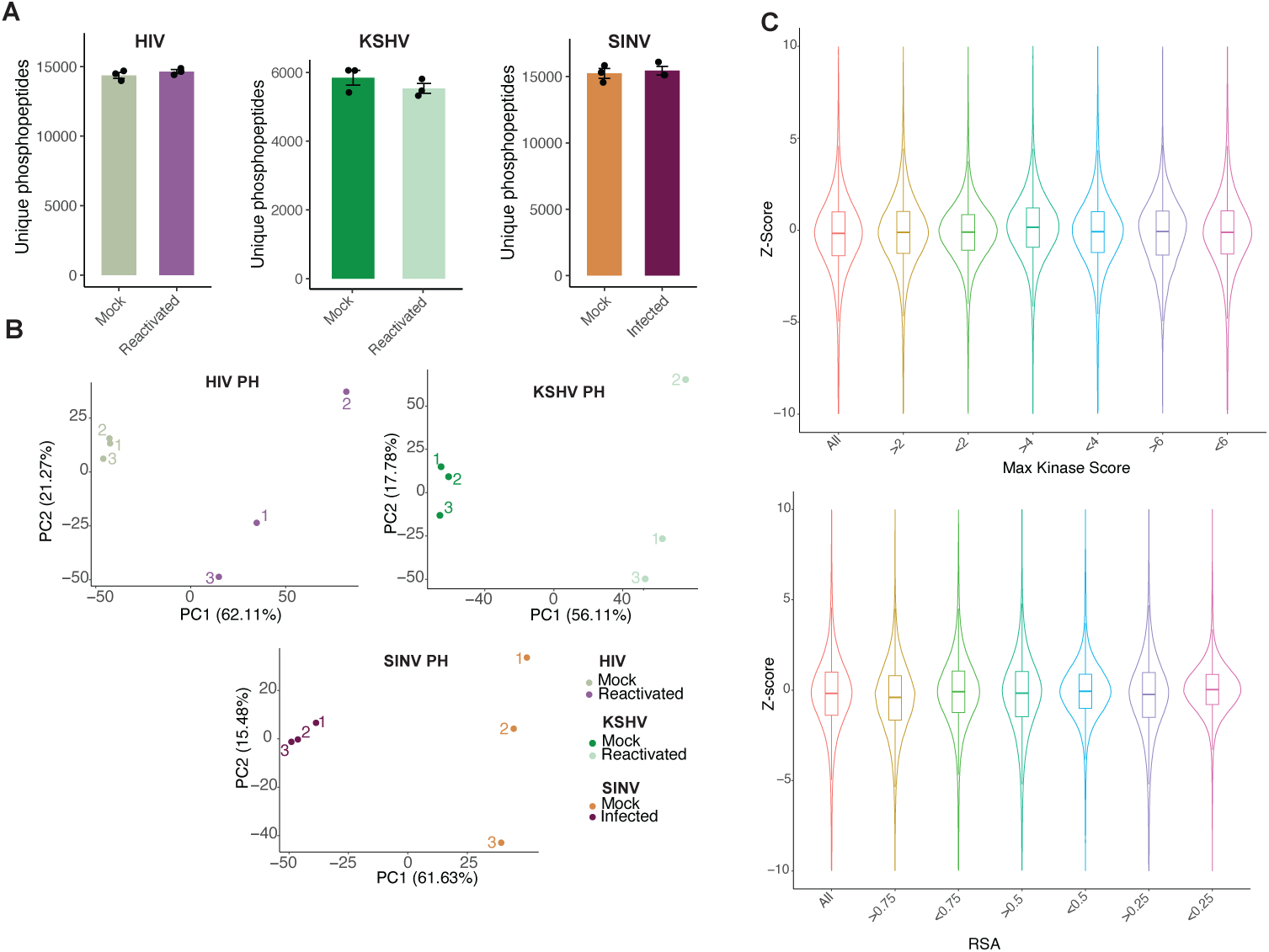
Quality control of phosphoproteomic samples and filtering threshold determination. (A) Number of unique phosphopeptides detected across three biological replicates for SINV infection in HEK293T cells, KSHV reactivation in B cells, and HIV reactivation in J-Lat 10.6 cells. (B) Principal component analysis (PCA) of phosphoproteomics data for SINV infection, and KSHV and HIV reactivation. (C) Comparison of different filtering thresholds using Z-score analyses for virus–human kinase comparisons. RSA and maximum kinase score were tested as filtering metrics with three different thresholds. Z-scores were computed per virus–kinase pair.

**Supplemental Figure 3.**
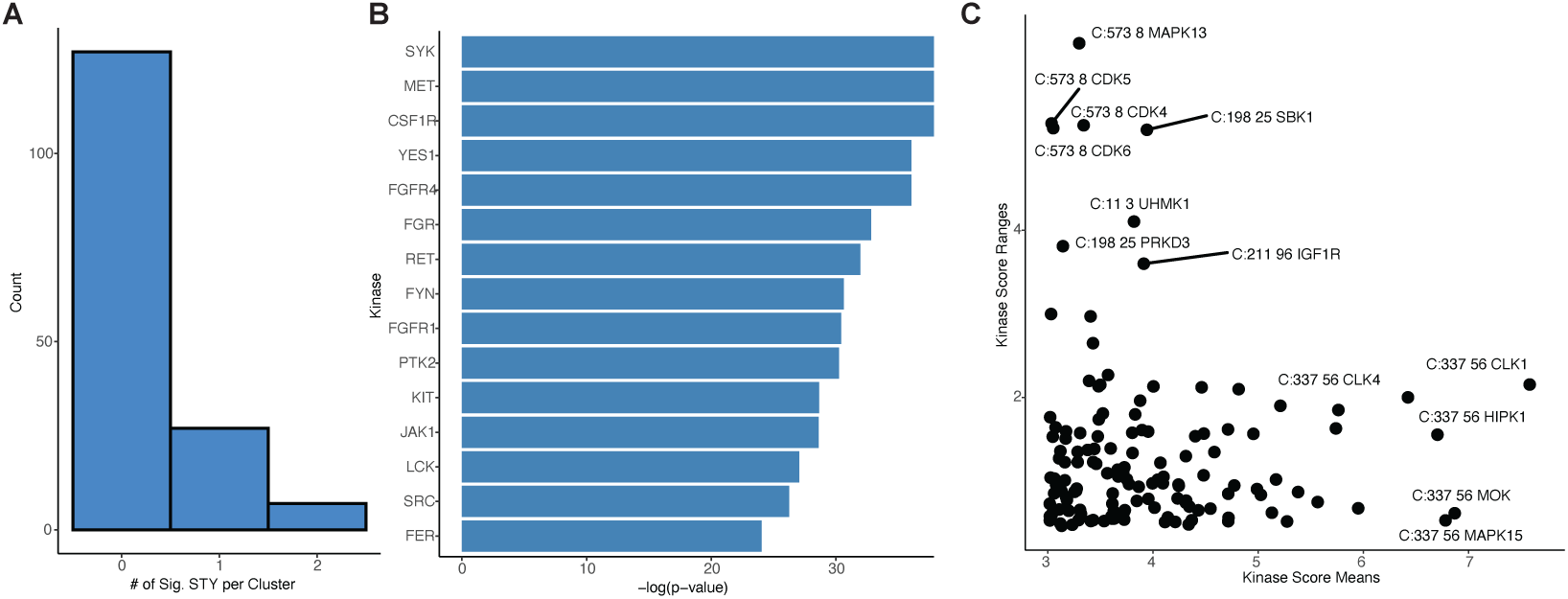
Extensions of positive selection analysis. (A) Number of serine, threonine, and tyrosine sites per cluster undergoing significant positive selection. (B) Top kinases predicted to have higher motif scores for sites under positive selection compared to non-significant sites. (C) Mean and range of kinase scores for sites with perfectly conserved phosphoacceptors and at least one positively selected residue in the adjacent motif. Values are space-separated and indicate cluster ID, amino acid position, and kinase.

**Supplemental Figure 4.**
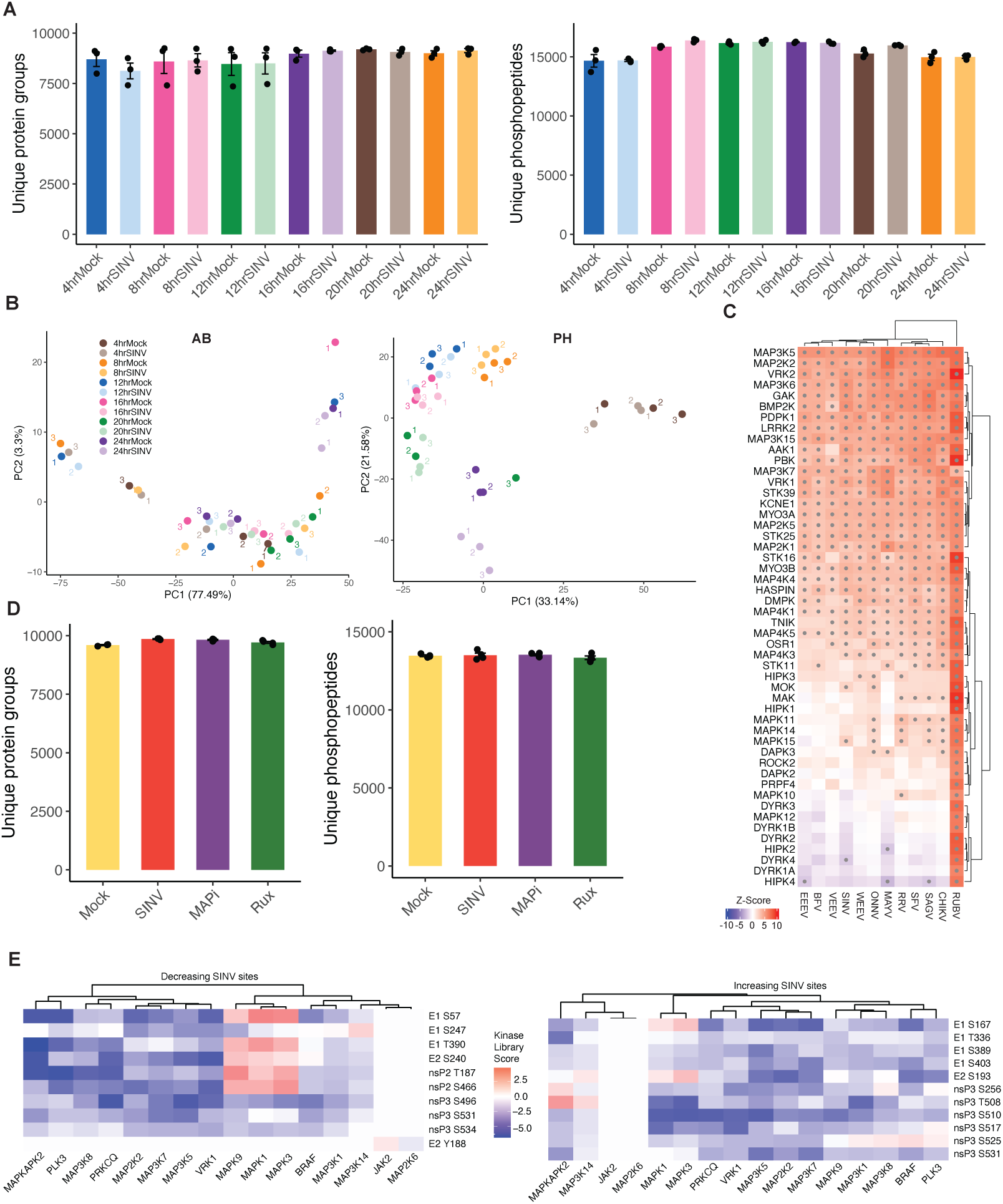
Sindbis virus time-course and inhibitor experimental quality control. (A) Number of unique protein groups (left) and unique phosphopeptides (right) across different conditions in the SINV time-course infection experiment. Each dot represents a biological replicate. (B) Principal component analysis (PCA) of total protein abundance data (left) and phosphoproteomics data (right) from the SINV time-course infection. (C) Z-scores for Togaviruses across various kinases within the stress-activated kinase family. (D) Number of unique protein groups (left) and unique phosphopeptides (right) across conditions in the SINV inhibitor experiment. Each dot represents a biological replicate. (E) Kinase Library scores for MAPK kinases corresponding to phosphosites that decrease (left) or increase (right) upon MAPK inhibitor treatment.

## DECLARATION OF INTERESTS

The authors declare no competing interests.

## ACKNOWLEDGEMENTS

We acknowledge the UCLA AIDS Institute, the James B. Pendleton Charitable Trust, and the McCarthy Family Foundation for their generous support of our research. We would like to acknowledge funding from the National Institutes of Health (NIH-NIGMS R35GM160071 to MB), a Center for AIDS Research (CFAR) Pilot Grant (1P30AI152501-01A1 to MB and OIF), and a Collaborative Development Award (CDA) from the HIV Accessory and Regulatory Complexes (HARC) center at the Quantitative Biosciences Institute (QBI) at UCSF (U54AI170792 to MB and OIF). Our trainees were generously supported by the Howard Hughes Medical Institute (HHMI) Gilliam Fellowship (#GT17454 to SKM), Molecular Biology Institute Whitcome Fellowship to SKM, the UCLA-Caltech Medical Scientist Training Program (NIH-NIGMS T32GM008042 to SFB), the David Geffen Medical Student Scholarship to SFB, the Cellular and Molecular Biology T32 at UCLA (NIH-NIGMS T32GM145388 to DMW), the Microbial Pathogenesis Training Grant (MPTG NIH-NIAID T32AI007323 to YD), the Graduate Research Fellowship Program (GRFP) from the National Science Foundation (DGE-2444110 to YD), and the Undergraduate Research Scholars Program Grant funded by the Van Trees (USRP, 2025-2026 to JN).

## DATA AVALIBILITY

Proteomics and large computational datasets used here were uploaded to the PRoteomics IDEntifications Database (PRIDE) at ebi.ac.uk/pride under project accession PXD070004. For reviewers, the token is “YeDRWD4jvJ74” and can be accessed using: Username: and Password: MVvirmphpmV1.

## SUPPLEMENTARY TABLES

**Table S1. Metadata for Human-Infecting Viruses**

Includes viral taxonomic classification, proteome composition, number of high-confidence kinase motifs, total STY sites, proteome size, and proportion of high-confidence motifs for all kinase families.

**Table S2. Z-Scores for Virus–Kinase Motif Enrichment**

Z-scores, p-values, and high-confidence motif counts for each virus–kinase pair, including significance marking based on applied thresholds.

**Table S3. GSOA Enrichment Results**

*Sheet 1:* GSOA enrichment for human-infecting viruses.

*Sheet 2:* GSOA enrichment for human-infecting viruses, grouped by viral families.

*Sheet 3:* GSOA enrichment for the Nomburg et al.^22^ dataset.

**Table S4. Positive Selection Analysis Results.**

*Sheet 1:* Viral protein clusters.

*Sheet 2:* MEME outputs per codon.

*Sheet 3:* GSOA enrichment for kinases preferentially phosphorylating sites undergoing positive selection.

**Table S5. Viral Protein Abundance and Phosphorylation in SINV Experiments**

*Sheet 1:* Summed viral abundance intensities over time during SINV infection. *Sheet 2:* Summed viral phosphorylation intensities over time during SINV infection.

*Sheet 3:* Summed viral abundance intensities in the inhibitor experiment.

*Sheet 4:* Summed viral phosphorylation levels in the inhibitor experiment.

**Table S6. Kinase Activity During SINV Time Course**

Calculated kinase activities based on phosphoproteomics data from the SINV time course.

## Notes

### Competing Interest Statement

The authors have declared no competing interest.

### Summary of Updates

A new Figure 4 has been added; the original Figure 4 and Figure 5 have been revised.

## REFERENCES

1. Duggal, N.K., and Emerman, M. (2012). Evolutionary conflicts between viruses and restriction factors shape immunity. Nat Rev Immunol 12, 687–695.

2. Huérfano, S., Šroller, V., Bruštíková, K., Horníková, L., and Forstová, J. (2022). The Interplay between Viruses and Host DNA Sensors. Viruses 14. 10.3390/v14040666.

3. Twarock, R., Towers, G.J., and Stockley, P.G. (2024). Molecular frustration: a hypothesis for regulation of viral infections. Trends Microbiol 32, 17–26.

4. Kumar, R., Mehta, D., Mishra, N., Nayak, D., and Sunil, S. (2020). Role of Host-Mediated Post-Translational Modifications (PTMs) in RNA Virus Pathogenesis. Int J Mol Sci 22. 10.3390/ijms22010323.

5. Ardito, F., Giuliani, M., Perrone, D., Troiano, G., and Lo Muzio, L. (2017). The crucial role of protein phosphorylation in cell signaling and its use as targeted therapy (Review). International Journal of Molecular Medicine 40, 271.

6. Keating, J.A., and Striker, R. (2012). Phosphorylation events during viral infections provide potential therapeutic targets. Rev Med Virol 22, 166–181.

7. Dey, S., and Mondal, A. (2024). Unveiling the role of host kinases at different steps of influenza A virus life cycle. J Virol 98, e0119223.

8. Ammosova, T., Berro, R., Jerebtsova, M., Jackson, A., Charles, S., Klase, Z., Southerland, W., Gordeuk, V.R., Kashanchi, F., and Nekhai, S. (2006). Phosphorylation of HIV-1 Tat by CDK2 in HIV-1 transcription. Retrovirology 3, 78.

9. Botova, M., Camacho-Zarco, A.R., Tognetti, J., Bessa, L.M., Guseva, S., Mikkola, E., Salvi, N., Maurin, D., Herrmann, T., and Blackledge, M. (2024). A specific phosphorylation-dependent conformational switch in SARS-CoV-2 nucleocapsid protein inhibits RNA binding. Sci Adv 10, eaax2323.

10. Patil, A., Anhlan, D., Ferrando, V., Mecate-Zambrano, A., Mellmann, A., Wixler, V., Boergeling, Y., and Ludwig, S. (2021). Phosphorylation of Influenza A Virus NS1 at Serine 205 Mediates Its Viral Polymerase-Enhancing Function. Journal of Virology 95, e02369–20.

11. Kitata, R.B., Choong, W.-K., Tsai, C.-F., Lin, P.-Y., Chen, B.-S., Chang, Y.-C., Nesvizhskii, A.I., Sung, T.-Y., and Chen, Y.-J. (2021). A data-independent acquisition-based global phosphoproteomics system enables deep profiling. Nat Commun 12, 2539.

12. Abramson, J., Adler, J., Dunger, J., Evans, R., Green, T., Pritzel, A., Ronneberger, O., Willmore, L., Ballard, A.J., Bambrick, J., et al. (2024). Addendum: Accurate structure prediction of biomolecular interactions with AlphaFold 3. Nature 636, E4.

13. Jeppesen, M., and André, I. (2023). Accurate prediction of protein assembly structure by combining AlphaFold and symmetrical docking. Nat Commun 14, 8283.

14. Johnson, J.L., Yaron, T.M., Huntsman, E.M., Kerelsky, A., Song, J., Regev, A., Lin, T.-Y., Liberatore, K., Cizin, D.M., Cohen, B.M., et al. (2023). An atlas of substrate specificities for the human serine/threonine kinome. Nature 613, 759–766.

15. Yaron-Barir, T.M., Joughin, B.A., Huntsman, E.M., Kerelsky, A., Cizin, D.M., Cohen, B.M., Regev, A., Song, J., Vasan, N., Lin, T.-Y., et al. (2024). The intrinsic substrate specificity of the human tyrosine kinome. Nature 629, 1174–1181.

16. Dochnal, S.A., Whitford, A.L., Francois, A.K., Krakowiak, P.A., Cuddy, S., and Cliffe, A.R. (2024). c-Jun signaling during initial HSV-1 infection modulates latency to enhance later reactivation in addition to directly promoting the progression to full reactivation. J Virol 98, e0176423.

17. Li, L., Li, P., Chen, A., Li, H., Liu, Z., Yu, L., and Hou, X. (2022). Quantitative proteomic analysis shows involvement of the p38 MAPK pathway in bovine parainfluenza virus type 3 replication. Virol J 19, 116.

18. Wang, J., Reuschel, E.L., Shackelford, J.M., Jeang, L., Shivers, D.K., Diehl, J.A., Yu, X.-F., and Finkel, T.H. (2011). HIV-1 Vif promotes the G₁-to S-phase cell-cycle transition. Blood 117, 1260–1269.

19. Slagle, B.L., and Bouchard, M.J. (2018). Role of HBx in hepatitis B virus persistence and its therapeutic implications. Curr Opin Virol 30, 32–38.

20. Aledavood, E., Selmi, B., Estarellas, C., Masetti, M., and Luque, F.J. (2021). From Acid Activation Mechanisms of Proton Conduction to Design of Inhibitors of the M2 Proton Channel of Influenza A Virus. Front Mol Biosci 8, 796229.

21. Murrell, Ben, Joel O. Wertheim, Sasha Moola, Thomas Weighill, Konrad Scheffler, and Sergei L. Kosakovsky Pond. 2012. “Detecting Individual Sites Subject to Episodic Diversifying Selection.” PLoS Genetics 8 (7): e1002764.

22. Nomburg, J., Doherty, E.E., Price, N., Bellieny-Rabelo, D., Zhu, Y.K., and Doudna, J.A. (2024). Birth of protein folds and functions in the virome. Nature 633, 710–717.

23. Murrell, B, et al. "Detecting individual sites subject to episodic diversifying selection." PLoS Genetics 8, e1002764 (2012).

24. Meyer, Austin G., and Claus O. Wilke. 2013. “Integrating Sequence Variation and Protein Structure to Identify Sites under Selection.” Molecular Biology and Evolution 30 (1): 36–44.

25. Taylor, Adam, Lara J. Herrero, Penny A. Rudd, and Suresh Mahalingam. 2015. “Mouse Models of Alphavirus-Induced Inflammatory Disease.” The Journal of General Virology 96 (Pt 2): 221–38.

26. Rincón, M., and Davis, R.J. (2009). Regulation of the immune response by stress-activated protein kinases. Immunological reviews 228. 10.1111/j.1600-065X.2008.00744.x.

27. Xie, J., Ajibade, A.O., Ye, F., Kuhne, K., and Gao, S.-J. (2008). Reactivation of Kaposi’s sarcoma-associated herpesvirus from latency requires MEK/ERK, JNK and p38 multiple mitogen-activated protein kinase pathways. Virology 371, 139–154.

28. Ma, Y., Kanakousaki, K., and Buttitta, L. (2015). How the cell cycle impacts chromatin architecture and influences cell fate. Front. Genet. 6, 126669.

29. Bagga, S., and Bouchard, M.J. (2014). Cell Cycle Regulation During Viral Infection. Cell Cycle Control 1170, 165.

30. Bolinger, C., and Boris-Lawrie, K. (2009). Mechanisms employed by retroviruses to exploit host factors for translational control of a complicated proteome. Retrovirology 6, 8.

31. Ciuffi, A. (2008). Mechanisms governing lentivirus integration site selection. Curr Gene Ther 8, 419–429.

32. Su, M.-G., and Lee, T.-Y. (2013). Incorporating substrate sequence motifs and spatial amino acid composition to identify kinase-specific phosphorylation sites on protein three-dimensional structures. BMC Bioinformatics 14 *Suppl 16*, S2.

33. Tien, Matthew Z., Austin G. Meyer, Dariya K. Sydykova, Stephanie J. Spielman, and Claus O. Wilke. 2013. “Maximum Allowed Solvent Accessibilites of Residues in Proteins.” PloS One 8 (11): e80635.

34. Chung, Cheng-Han, Anthony R. Mele, Alexander G. Allen, Robert Costello, Will Dampier, Michael R. Nonnemacher, and Brian Wigdahl. 2020. “Integrated Human Immunodeficiency Virus Type 1 Sequence in J-Lat 10.6.” Microbiology Resource Announcements 9 (18). 10.1128/MRA.00179-20.

35. Karline Soetaert, Thomas Petzoldt, R. Woodrow Setzer (2010). “Solving Differential Equations in R: Package deSolve.” Journal of Statistical Software, 33(9), 1–25. doi:10.18637/jss.v033.i09.

36. Türei, Dénes, Alberto Valdeolivas, Lejla Gul, Nicolàs Palacio-Escat, Michal Klein, Olga Ivanova, Márton Ölbei, et al. 2021. “Integrated Intra- and Intercellular Signaling Knowledge for Multicellular Omics Analysis.” Molecular Systems Biology 17 (3): e9923.

37 Casado, Pedro, Juan-Carlos Rodriguez-Prados, Sabina C. Cosulich, Sylvie Guichard, Bart Vanhaesebroeck, Simon Joel, and Pedro R. Cutillas. 2013. “Kinase-Substrate Enrichment Analysis Provides Insights into the Heterogeneity of Signaling Pathway Activation in Leukemia Cells.” Science Signaling 6 (268): rs6.

38. Hernandez-Armenta, Claudia, David Ochoa, Emanuel Gonçalves, Julio Saez-Rodriguez, and Pedro Beltrao. 2017. “Benchmarking Substrate-Based Kinase Activity Inference Using Phosphoproteomic Data.” Bioinformatics (Oxford, England) 33 (12): 1845–51.

39. Müller-Dott, Sophia, Eric J. Jaehnig, Khoi Pham Munchic, Wen Jiang, Tomer M. Yaron-Barir, Sara R. Savage, Martin Garrido-Rodriguez, et al. 2025. “Comprehensive Evaluation of Phosphoproteomic-Based Kinase Activity Inference.” Nature Communications 16 (1): 4771.

